# Revealing the neural networks that extract conceptual gestalts from continuously evolving or changing semantic contexts

**DOI:** 10.1101/666370

**Authors:** Francesca M. Branzi, Gina F. Humphreys, Paul Hoffman, Matthew A. Lambon Ralph

## Abstract

Reading a book, understanding the news reports or any other behaviour involving the processing of meaningful stimuli requires the semantic system to have two main features: being active during an extended period of time and flexibly adapting the internal representation according to the changing environment. Despite being key features of many everyday tasks, formation and updating of the semantic “gestalt” are still poorly understood. In this fMRI study we used naturalistic stimuli and task manipulations to identify the neural network that forms and updates conceptual gestalts during time-extended integration of meaningful stimuli. Univariate and multivariate techniques allowed at drawing a distinction between networks that are crucial for the formation of a semantic gestalt (meaning integration) and those that instead are important for linking incoming cues about the current context (e.g., time, space cues) into a schema representation. Specifically, we revealed that time-extended formation of the conceptual gestalt was reflected in the neurocomputations of the anterior temporal lobe accompanied by multi-demand areas and hippocampus, with a key role of brain structures in the right hemisphere. This “semantic gestalt network” was strongly recruited when an update of the current semantic representation was required during narrative processing. A distinct fronto-parietal network, instead, was recruited for context integration, independently from the meaning associations between words (semantic coherence). Finally, in contrast with accounts positing that the default-mode-network (DMN) may have a crucial role in semantic cognition, our findings revealed that DMN activity was sensitive to task difficulty, but not to semantic integration. The implications of these findings for neurocognitive models of semantic cognition and the literature on narrative processing are discussed.

**Highlights:**

- fMRI revealed areas and networks for semantic integration during narrative reading
- ATL has a key role in the formation of the conceptual gestalt
- IFG, pMTG and dAG reflect the update of the conceptual gestalt
- Left AG (Mid-PGp) has a key role in context integration

## 1. Introduction

Successful time-extended semantic cognition (e.g., understanding the news reports on the radio or sequential paragraphs while reading a book) relies on the ability of the semantic system to integrate information over time to build meaningful representations of the evolving world around us. Although semantic integration is often error-free and apparently effortless, the cognitive challenges are non-trivial. Thus, the semantic system is required over an extended-period of time to build a continuously evolving conceptual gestalt from the torrent of words and non-verbal stimuli yet, when the context or situation changes (which is often not overtly signalled but has to be inferred), the semantic system has to update or reset and build a new conceptual gestalt afresh.

Despite being a core, everyday function of semantic cognition, the neural foundations of conceptual gestalt formation and update are still uncharacterised. In fact, a handful of studies have utilised word pairs to explore the formation of meaning at the noun-phrase level (Price et al., 2015; Price et al., 2016) without addressing the time-extended demands posed by everyday semantic cognition. Other studies, instead, have measured multi-item semantic combinations without task manipulations that distinguish between brain structures involved in semantic integration (or formation of the semantic gestalt) from those involved in extra-semantic neurocomputations (e.g., working memory (WM), attentional control, etc.: e.g., (Hasson et al., 2008; Lerner et al., 2011; Simony et al., 2016; Tylen et al., 2015).

Accordingly, in this functional magnetic resonance imaging (fMRI) study we established the brain regions and networks that support time-extended semantic integration (formation of the semantic gestalt), as well as those that are sensitive to changes in semantic context (update of the semantic gestalt). Importantly, in doing so we provide some timely evidence that bridges the existing work on narratives (Hasson et al., 2008; Lerner et al., 2011; Simony et al., 2016; Tylen et al., 2015) with the broader neurocognitive theories of semantic cognition (Lambon Ralph et al., 2017).

The neural foundations of conceptual gestalt formation and update were addressed through a straightforward fMRI experimental design. First, we used combinations of short paragraphs to reflect the time-extended demands posed by everyday semantic cognition and the challenges posed by the formation of larger conceptual gestalts. Second, we adopted a key manipulation to determine the neural networks involved in the formation of the conceptual gestalt and to differentiate them from brain structures supporting extra-semantic control mechanisms. Thus, participants were asked to read short narratives composed of two phases (context and target). For each narrative, the same second paragraph (target) was preceded by different types of context: (i) a highly congruent context (HC) which maximised the information contained in a single coherent semantic gestalt; (ii) a low-congruent (LC) paragraph with a divergent meaning, thus testing the semantic system’s ability to update the information presented context (see Table S1); and (iii) a no context control (NC) in which the same target was preceded by a number reading task (thus requiring a more substantial shift from a non-semantic to semantic task and at the same time preventing context integration). The resultant fMRI data were analysed only for the identical target paragraph (thus ensuring that any observed differences must reflect the influence of the preceding contexts) that was compared across conditions to determine the neural foundations of formation (HC&LC>NC) and update (LC>HC) of the semantic gestalt. To establish the key brain areas and networks, we used both univariate and multivariate (independent component analysis; ICA) analyses. Particularly, we combined a whole-brain data-driven analytic approach, here particularly appropriate given the complexity of the investigated phenomena, with an independent ROI analysis approach, to link the present work to previous investigations on semantic cognition (Lambon Ralph et al., 2017).

According to previous studies, we expected to observe brain areas already implicated in semantic cognition as well as additional networks that might reflect the demands posed by formation and updating of conceptual contexts. A large body of cognitive and clinical neuroscience research, based primarily on the processing and representation of single concepts, has identified two interactive neural networks (Lambon Ralph et al., 2017). The first builds coherent, generalizable, multimodal conceptual representations through the interaction of an anterior temporal lobe (ATL) hub with a distributed set of secondary association cortices (Lambon Ralph et al., 2010; Patterson et al., 2007; Rogers et al., 2004). The computational models of this network (Chen et al., 2017b; Hoffman et al., 2018; Rogers et al., 2004) have always emphasised that the ATL hub allows information to be combined into coherent concepts from across verbal and nonverbal sources (words, sounds, vision, etc.) and also over time. Accordingly, it seems entirely possible that this network would be important for the formation of the conceptual gestalts conveyed in the narratives (i.e., HC&LC>NC). Indeed, a handful of previous studies have found evidence for a combinatorial semantic process in the superior ATL (Brennan and Pylkkanen, 2012; Humphries et al., 2006; Maguire et al., 1999; Noppeney and Price, 2004; Pallier et al., 2011; Vandenberghe et al., 2002).

The second established semantic network reflects the need to shape and mould semantic information to align with changing tasks and contexts demands (Jefferies and Lambon Ralph, 2006; Noonan et al., 2013). This executive semantic network is comprised of prefrontal, posterior middle temporal and intraparietal sulcus areas (Humphreys and Lambon Ralph, 2015; Noonan et al., 2013). It seems likely that this second network will be important in processing the meaning of the narratives, especially when there is a shift in the context (updating of the semantic gestalt, i.e., LC>HC).

Given the cognitive requirements posed by the formation of meaning across a complete narrative, additional brain regions and networks are likely to be engaged. ICA provides a data-driven approach for identifying independent spatiotemporal functional networks, which can then be tested for their sensitivity to the experimental conditions included in the fMRI task. Thus, ICA is particularly suitable for revealing which brain structures are recruited simultaneously for semantic integration. Previous studies have shown that task-active and resting-state fMRI reveal multiple spatiotemporal networks including a semantic-language network (SLN), an executive control network (ECN), a default-mode network (DMN), etc. (Beckmann et al., 2005; Geranmayeh et al., 2014; Seeley et al., 2007; Wirth et al., 2011). Some of these networks overlap with the hypothesised neural networks for semantic representation and control (see above). Thus, consistent with the literature on narratives (Hasson et al., 2008; Lerner et al., 2011; Price, 2012; Simony et al., 2016; Vigneau et al., 2006; Xu et al., 2005) and the hub and spokes model (Lambon Ralph et al., 2017), we expected some of these to be engaged by the task and be sensitive to the task manipulations. In detail, the networks associated with dorsal executive and semantic control should be critically important for update of the semantic gestalt (i.e., LC>HC), as they have been shown to engage whenever semantic processing is demanding (Hoffman et al., 2015; Noonan et al., 2013; Tylen et al., 2015) or after changes of non-semantic rules (Vatansever et al., 2017b).

Furthermore, the DMN – a network comprising midline brain regions and the angular gyrus (AG) – which is often anticorrelated with the dorsal executive network (Fox et al., 2005; Humphreys and Lambon Ralph, 2017; Spreng et al., 2010; Vincent et al., 2008, but see Dixon et al., 2017) – might also be a crucial network for the formation of conceptual gestalts. Its exact role, though, is hard to predict from the current literature on the DMN. On the one hand, some researchers have suggested that the DMN may support processes for semantic integration (Hasson et al., 2008; Lerner et al., 2011; Simony et al., 2016; Tylen et al., 2015; Wirth et al., 2011), which would align with the seminal work of Binder that ‘rest’ involves considerable spontaneous semantic-language activities (Binder et al., 2009). More recent evidence indicates that this hypothesis holds for nodes within the semantic network, such as the ATL, but the AG and other aspects of the DMN are deactivated by semantic and non-semantic tasks alike (Humphreys et al., 2015; Humphreys and Lambon Ralph, 2017). According to the proposal that DMN and AG are actively involved in the formation of semantic gestalt, this network should be positively engaged during semantic processing (i.e., in all conditions), in a way proportional to the amount (HC&LC>NC) and congruency (HC>LC) of the semantic information to be integrated. On the other hand, explorations of episodic memory have implicated the AG and DMN in vivid episodic retrieval and buffering (Rugg and Vilberg, 2013; van der Linden et al., 2017; Vatansever et al., 2017a; Vilberg and Rugg, 2008). Accordingly, it is possible that this buffering mechanism might be engaged by the time-extended narratives whilst participants build up a mental model for the story based on the pre-existing context/schema (Ranganath and Ritchey, 2012) (i.e., positively engaged during HC and LC conditions only), predicting greatest activation for the consistent context condition (i.e., HC>LC) (van der Linden et al., 2017; Vatansever et al., 2017b). Interestingly, this hypothesis would be also in accord with evidence showing that the DMN is modulated by time-extended semantic integration (Hasson et al., 2008; Simony et al., 2016). Finally, an alternative hypothesis with an opposing prediction arises from recent studies that suggest that the DMN is engaged when switching between activities (Crittenden et al., 2015; Smith et al., 2018). If correct, then the DMN should be positively engaged during switches of task or semantic contexts (i.e., during NC and LC conditions, respectively) and particularly when switching from a number to language activity (i.e., NC>LC), but disengaged when no switch is perceived (i.e., HC conditions).

## 2. Materials and Methods

### 2.1. Participants

Twenty-four volunteers took part in the study (average age=22, SD=2; N female=18). All participants were native English speakers with no history of neurological or psychiatric disorders and normal or corrected-to-normal vision. As a result of technical issues during the scanning session, only data from 22 participants (average age=22 years, SD=2; N female=18) were usable for fMRI data analyses. The work described has been carried out in accordance with The Code of Ethics of the World Medical Association (Declaration of Helsinki) for experiments involving humans. Furthermore, all participants gave written informed consent and the study was approved by the local ethics board.

### 2.2. Stimuli

A total of 40 narrative pairs, each one composed by two paragraphs, were created for the experimental study. For each narrative pair, the same second paragraph (target) was preceded by different first paragraphs (contexts) that could be either high-congruent (i.e., HC) or low-congruent (i.e., LC) with the target in terms of meaning. Both HC and LC context paragraphs could be integrated with the targets, though a reworking of the evolving semantic context was required after LC contexts only, because of a shift in the semantic context (see Table S1 for the complete list of the stimuli). Homonym words (e.g., bank) were employed in order to determine the exact point in the paragraph in which the shift in the semantic context should have been experienced.

To ensure that HC and LC conditions differed in respect to semantic associative strength between contexts and targets, we quantified in different ways semantic relatedness between the contexts and targets for both HC and LC conditions. First, we employed Latent Semantic Analysis (LSA) (Hoffman, 2019; Hoffman et al., 2013; Landauer and Dumais, 1997), a method to measure the semantic relationship between words based on the degree to which they are used in similar linguistic contexts. Hence, for each narrative pair, pairwise LSA values were calculated for contexts and targets and then averaged within both. As result, an LSA value reflecting the associative strength between the context and target was obtained for both conditions. Results from LSA confirmed that semantic associative strength between the (same) target and the context was higher for HC (average score=0.41, SD=0.16) than LC conditions (average score=0.23, SD=0.1) [t (78)=−5.996, p<0.001].

Second, we asked to a group of independent participants to indicate how much contexts and targets were perceived as being semantically related (0 to 5 scale). The results of this pre-experimental rating confirmed that HC (average score=4.4, SD=0.4) and LC (average score=2.3, SD=0.4) conditions were different [t(9)=−10.626, p<0.001]. Moreover, to ensure that participants could perceive the shift of semantic context during the study, at end of each narrative the question “Was there any change of semantic context between part1 and part2?” was posed. Only pairs of narratives on which at least the 90% of participants responded correctly to the questions were employed in the study.

Finally, another condition was included in the design in order to measure the semantic integration processes in general. Precisely, in the NC condition the target (the same as in HC and LC conditions) was preceded by a string of numbers that could include from one to four-digit numbers.

### 2.3. Task procedures

There were 40 items per condition presented using an event-related design with the most efficient ordering of events determined using Optseq (http://www.freesurfer.net/optseq). Rest time was intermixed between trials and varied between 2 and 12 seconds (s) (average=3.7, SD=2.8) during which a red fixation cross was presented. The red colour was used in order to mark the end of each trial (each narrative composed by a context and a target). A black fixation cross was presented between contexts and targets and its duration varied between 0 and 6s (average=3, SD=1.6). Each context paragraph was presented for 9s followed by the target for 6s.

Participants were asked to read silently both contexts (verbal material and numbers) and targets (only verbal material). Our volunteers were instructed to press a button when arriving to the end of each paragraph (for both contexts and targets). The instruction emphasized speed, but also the need to understand the meaning of verbal contexts and targets, since at the end of some of the trials participants would have been asked to answer to some questions on the content of the narratives. We specified that in order to perform this task it would have been necessary to integrate the meaning between contexts and targets. Hence, following 13% of the trials a comprehension task was presented to ensure that participants were engaged in the task. When this happened, the target item was followed by a question displayed on the screen for 6s at which participants were required to provide a response (true/false) via button press. A fixation cross between the target and the question was presented during a time that varied between 0 and 6s (average=3.5, SD=2.2). Before starting the experimental study, all participants were given written instructions. Then they underwent to a practice session with few trials in order to allow them to familiarise with the task. The stimuli used in the practice session were different from those used in the experimental study.

### 2.4. Task acquisition parameters

Images were acquired using a 3T Philips Achieva scanner using a dual gradient-echo sequence, which is known to have improved signal relative to conventional techniques, especially in areas associated with signal loss (Halai et al., 2014). Thus, 31 axial slices were collected using a TR=2s, TE=12 and 35 milliseconds (ms), flip angle=95°, 80 × 79 matrix, with resolution 3 × 3mm, slice thickness 4mm. For each participant, 1492 volumes were acquired in total, collected in four runs of 746s each.

### 2.5. Data analysis

#### Behavioural data analyses

Behavioural analyses were performed on RTs and the percentage of given responses. Two separate repeated-measures analyses of variance (ANOVAs), one for RTs and the other for percentage of given responses, with “Condition” as within-subjects factor with three levels (NC, LC and HC conditions) were conducted. Bonferroni correction for multiple comparisons was applied to assess statistically significant effects.

#### fMRI data analyses

##### Preprocessing

The dual-echo images were averaged. Data was analysed using SPM8. After motion-correction images were co-registered to the participant’s T1 image. Spatial normalisation into MNI space was computed using DARTEL (Ashburner, 2007), and the functional images were resampled to a 3 × 3 × 3mm voxel size and smoothed with an 8mm FWHM Gaussian kernel.

#### General Linear Modelling (GLM)

The data was filtered using a high-pass filter with a cut-off of 128s and then analysed using a GLM. At the individual subject level, each condition was modelled with a separate regressor (Target NC, Target LC and Target HC) with time derivatives added, and events were convolved with the canonical hemodynamic response function, starting from the onset of the target paragraph. The number reading paragraph condition (Context NC) was also modelled as a regressor of interest in order to have an active baseline against which to compare the semantic tasks. Also the other context paragraphs and comprehension task were modelled as separate regressors of no interest. Each condition was modelled as a single event. Motion parameters were entered into the model as covariates of no interest.

#### The Semantic Network

To identify the brain areas involved in semantic processing during the narrative reading task, we assessed the whole-brain contrast of semantic target conditions (NC, LC, HC collapsed) against rest and against the number reading task (Context NC condition). All the contrasts were corrected for multiple comparisons with a voxel-wise false discovery rate (FDR) threshold set at *q*<0.05 (Benjamini and Hochberg, 1995) and a contiguity threshold ≥30 voxels.

Having identified the semantic network, we conducted region of interest (ROI) analyses to assess the functional contribution of key semantic areas in respect to our task manipulations. All the ROI coordinates were independently derived from the literature (Table S2). Regarding the parietal ROIs, we investigated the functional role of three different portions of the AG (Humphreys et al., 2019). We employed 10mm spheres for these analyses. Repeated-measures ANOVAs with “Condition” as a within-subjects factor with three levels (NC, LC and HC targets) were conducted to for temporal, semantic control and AG ROIs. Bonferroni correction for multiple comparisons was applied to assess statistically significant effects.

#### Whole-brain univariate analyses of the differences between experimental conditions

##### Semantic integration effect (or formation of the conceptual gestalt

To investigate the semantic integration effect the contrasts LC>NC and HC>NC were computed via whole-brain analysis. All the contrasts were corrected for multiple comparisons with a voxel-wise FDR threshold set at *q*<0.05 and a contiguity threshold ≥30 voxels.

##### Shift of semantic context (or update of the conceptual gestalt) and shift of task context

The shift of semantic context effect was established by running whole-brain analysis for the contrast LC>HC. To reveal brain regions that are important for task switches into language from a non-language task (i.e., shift of task context), we also conducted a NC>HC whole-brain contrast. Also these contrasts were corrected for multiple comparisons with a voxel-wise FDR correction threshold set at *q*<0.05 and a contiguity threshold ≥30 voxels.

Importantly, whole-brain contrast analyses alone do not inform on whether the observed differential activation is originated by task-positive or task-negative activation disparities. Hence, the precise contribution of each area was established by conducting ROI analysis (via repeated-measures ANOVA) on a set of key regions revealed by the whole-brain univariate analyses above. In this analysis we opted for 8mm spheres in order to restrict our ROIs only to voxels found to be significantly activated in the univariate contrasts.

#### Task group spatial ICA

Spatial ICA applied to fMRI data identifies temporally coherent networks by estimating maximally independent spatial sources, referred to as spatial maps (SMs), from their linearly mixed fMRI signals, referred to as time courses (TCs). The pre-processed fMRI data was analysed in a group spatial ICA using the GIFT toolbox (http://mialab.mrn.org/software/gift) (Calhoun et al., 2001) to decompose the data into its components. GIFT was used to concatenate the subjects’ data, and reduce the aggregated data set to the estimated number of dimensions using principal component analysis (PCA), followed by an ICA analysis using the Infomax algorithm (Bell and Sejnowski, 1995). Subject-specific SMs and TCs were estimated using GICA back-reconstruction method based on PCA compression and projection (Calhoun et al., 2001).

The number of independent components (ICs) estimated within the data was 38. The estimation was achieved by using the Minimum Description Length criteria, first per each individual data-set and then computing the group mean. The obtained 38 ICs were inspected in order to exclude from the analysis artefactual and noise-related components. Similar to previous studies (Geranmayeh et al., 2014; Griffanti et al., 2017) the criterion for assigning components as artefact was based on the SMs attained as a result of the one sample t-tests (threshold for voxel-wise significance was set at p<0.05, corrected for family-wise error (FWE), and a contiguity threshold ≥30 voxels). The SMs were visually compared with the SPM grey matter template. Only components that had the majority of activity within the grey matter were selected (N=22).

##### Establishing task-related FNs

The 22 ICs were labelled according to the resting state networks template provided in the GIFT toolbox. Then, a multiple regression analysis (implemented as “temporal sorting” function in GIFT) between IC and task model’s TCs for each participant was conducted and allowed to identify the ICs related to target conditions (task-related FNs). For that, for each participant the design matrix used for the GLM analysis, where rest periods were modelled implicitly as task baseline, was employed. For each IC, the multiple regression analysis generated 3 beta weight values (one for each condition NC, LC, and HC) that were averaged across runs and participants. Beta weight values represent the correlations between TCs of the ICs and the canonical hemodynamic response model for each task condition. These values are thought to reflect engagement of the functional networks (FNs) during specific task conditions (Xu et al., 2013).

Once extracted the beta weights for each IC associated with each condition, task-relatedness for each IC was assessed by testing group means of averaged beta weights for each task-condition against zero (one-sample t-tests, p<0.05). Hence, a positive/negative beta weight value significantly different from zero indicates increase/decrease in activity of the IC during a specific task condition relative to the baseline condition (i.e., rest). Once established the task-related FNs, for each FN a repeated-measures ANOVA was used to assess the main differences between beta weights across different task-conditions. Bonferroni correction for multiple comparisons was applied to assess statistically significant effects.

##### Functional network connectivity (FNC) analysis

To investigate the relationship between task-related FNs, we conducted a FNC analysis using the Mancovan toolbox in GIFT. Hence, FNC was estimated as the Pearson’s correlation coefficient between pairs of TCs (Jafri et al., 2008). Subject specific TCs were detrended and despiked based on the median absolute deviation as implemented in 3dDespike (http://afni.nimh.nih.gov/), then filtered using a fifth-order Butterworth low-pass filter with a high frequency cutoff of 0.15 Hz. Pairwise correlations were computed between TCs, resulting in a symmetric c1 × c1 correlation matrix for each subject. For all FNC analyses, correlations were transformed to z-scores using Fisher’s transformation, z=atanh(k), where k is the correlation between two component TCs. One sample t-tests (corrected for multiple comparisons at α=0.01 significance level using FDR) were conducted on task-related FNs in order to reveal the significance of pairwise correlations.

**Data availability statement.** The data will be made available at http://www.mrc-cbu.cam.ac.uk/publications/opendata/.

## 3. Results

### 3.1. Behavioural results

The reading times (RTs) showed the expected behavioural effect of semantic coherence. Thus performance differed across experimental conditions [F(2,46)=7.109, p=0.002, *η*p^2^=0.236], with reading speed in the NC and LC conditions being slower than in the HC condition (p=0.008 and p=0.019, respectively). There was no significant difference for reading the target sentence after the NC or LC contexts (p>0.999).

The percentage of given responses was very high and similar across all experimental conditions [F(2,46)=2.521, p=0.091, *η*p^2^=0.099], with a trend towards significance for percentage of given responses in the HC condition being higher than in the NC condition (p=0.098) (Figure 1). As mentioned in the Materials and Methods section, two participants were excluded from the fMRI analyses. Hence, we also ran behavioural analyses for these 22 participants. The results remained unchanged from those reported above for both RTs [F(2,42)=5.491, p=0.008, *η*p^2^=0.207; NC>HC (p=0.02); LC>HC (p=0.05); NC=LC (p>0.999)] and percentage of given responses [F(2,42)=2.841, p=0.07, *η*p^2^=0.119; NC<HC (p=0.077); LC=HC (p>0.999); NC=LC (p=0.524)].

**Figure 1.**
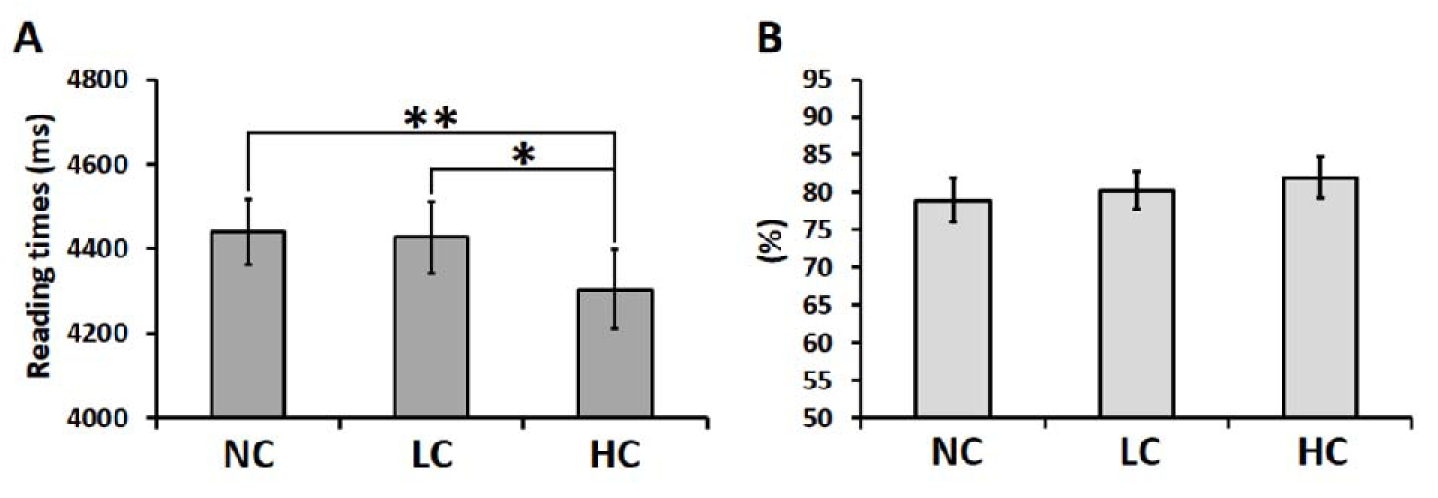
Behavioural results for **(A)** reading times and **(B)** percentage of given responses for NC, LC and HC target conditions. Error bars correspond to Standard Error (SE).

### 3.2. fMRI results

#### 3.2.1. GLM results

##### The Semantic Network

Whole-brain univariate analysis revealed that semantic reading tasks (target conditions) against rest or non-semantic reading tasks (number reading context) recruit a similar network of brain areas (Figure 2 and Table S3). This network includes frontal, temporal, and parietal brain areas, previously identified as key regions supporting semantic cognition (Lambon Ralph et al., 2017). Furthermore, time-extended reading tasks recruit extensively also the right hemisphere and other areas, normally deactivated during semantic tasks (e.g., hippocampus, precuneus, and the mid-PGp portion of the left AG) (Humphreys et al., 2015; Humphreys and Lambon Ralph, 2015, 2017).

**Figure 2.**
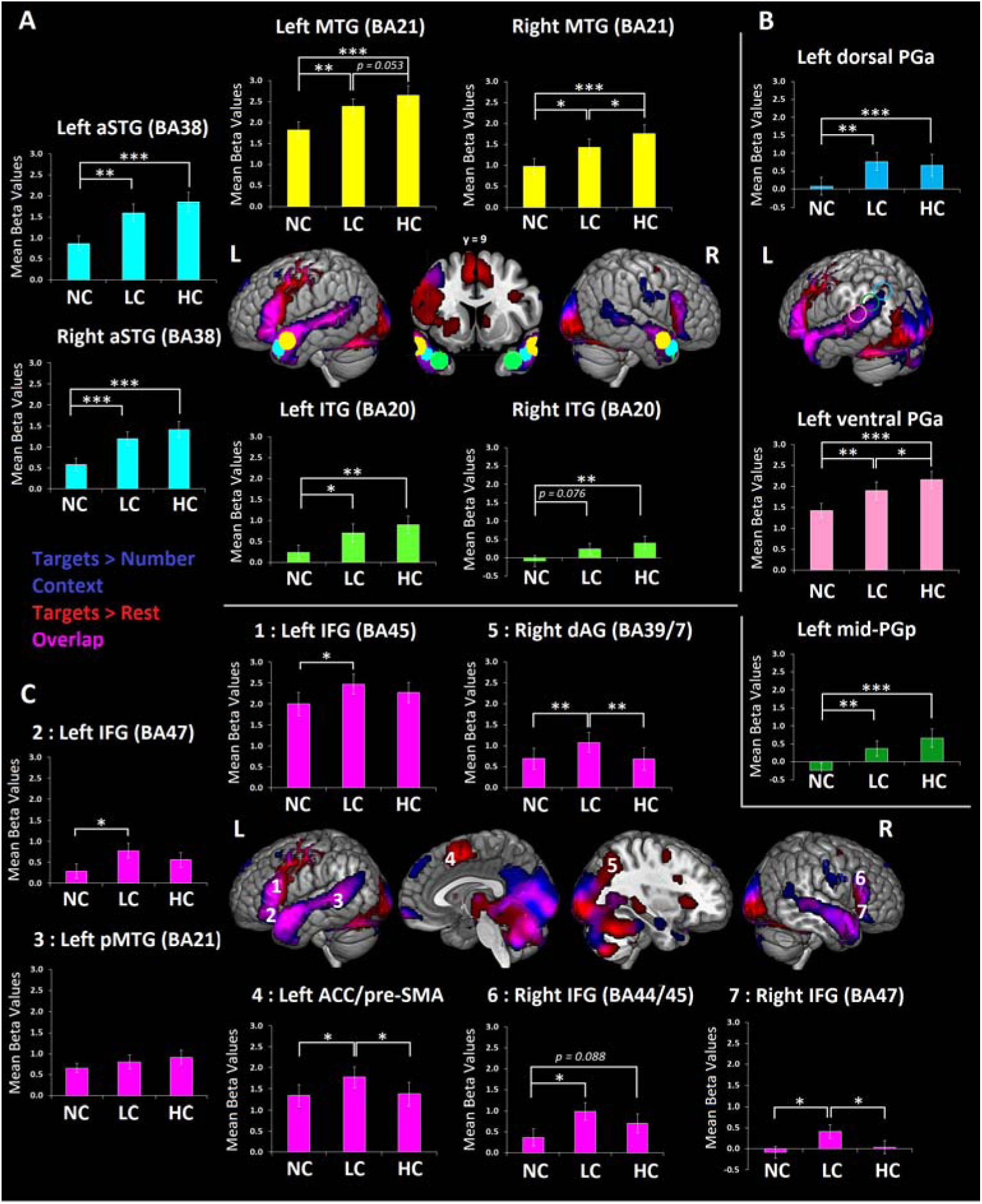
GLM results for target conditions against passive and active baselines and for different ROIs (derived from the literature) including **(A)** the ATL **(B)** the left AG and **(C)** the semantic control network. Mean beta values for each task condition were compared against rest. Error bars correspond to SE.

To determine which parts of the semantic network were sensitive to task manipulations, a set of independently-derived ROIs were employed (see Table S2). These ROIs included ventral and dorsal portions of the ATL, left AG, posterior middle temporal gyrus (pMTG), inferior frontal gyrus (IFG), and anterior cingulate cortex/pre-supplementary motor area (ACC/pre-SMA) (Figure 2).

###### Temporal lobe ROIs

We tested whether neural responses for each task-condition (NC, LC and HC) were statistically different across the three ATL regions (MTG, STG and ITG) and hemispheres (left and right), by conducting a repeated-measures ANOVA with “Region” (MTG, STG and ITG), “Hemisphere” (left and right) and “Condition” (NC, LC and HC) as within-subjects factors. A significant interaction between “Region” and “Condition” suggested that neural responses in the temporal ROIs were differently modulated by task conditions [F(4,84)=7.472, p<0.001, *η*p^2^=0.262]. Bonferroni-corrected post-hoc t-tests revealed that anterior MTG (∼BA21) showed significantly increased responses for conditions with integration (LC and HC conditions) as compared to NC condition (p=0.005 and p<0.001, respectively), and particularly for coherent semantic endings (HC>LC, p=0.016). Second, the anterior superior temporal gyrus (aSTG∼BA38) showed increased neural responses for paragraphs preceded by a contextual support (LC and HC conditions) as compared with paragraphs without contextual integration (NC condition) (p=0.001 and p<0.001, respectively). However, no significant difference between was observed between LC and HC conditions (p=0.1). Third, the inferior temporal gyrus (ITG∼BA20) showed sensitivity to integration of the contextual support similarly as the aSTG. In fact, neural activity in this region was increased for LC and HC conditions as compared with the NC condition (p=0.043 and p=0.001, respectively), but no significant difference was observed between LC and HC conditions (p=0.184). Importantly, ITG was not positively engaged (compared to rest) across all experimental conditions. Precisely, negligible responses were observed for NC condition [left ITG: t(21)=1.36, p=0.188; Right ITG: t(21)=−0.605, p=0.552] and for LC condition in the right hemisphere [t(21)=1.537, p=0.139], suggesting that this area was mostly engaged during integration of coherent semantic information (HC condition). Finally, the “Region”, “Condition” and “Hemisphere” triple interaction was not significant, suggesting that the condition-specific modulations observed in MTG, aSTG and ITG were similar in the right and left hemispheres [F(4,84)=0.466, p=0.760, *η*p^2^=0.022].

###### Left AG ROIs

We tested whether neural responses for each task-condition (NC, LC and HC) were statistically different across the three different left AG spheres. Hence, we conducted a repeated-measures ANOVA with “Region” (mid-PGp, ventral PGa and dorsal PGa) and “Condition” (NC, LC and HC) as within-subjects factors. A significant interaction between “Region” and “Condition” confirmed a different profile of engagement of the three different AG regions during the different task conditions [F (4,84)=5.668, p<0.001, *η*p^2^ = 0.213]. Bonferroni-corrected post-hoc t-tests revealed that the mid-PGp showed increased neural responses for paragraphs preceded by a contextual support (LC and HC conditions) as compared with paragraphs without contextual integration (NC condition) (p=0.009 and p<0.001, respectively). However, no significant difference was observed between LC and HC conditions (p=0.16). The ventral PGa also showed increased neural responses for LC and HC conditions as compared with paragraphs without contextual integration (NC condition) (p=0.006 and p<0.001, respectively), and particularly for coherent semantic endings (HC>LC, p=0.015). Finally, the dorsal PGa showed a similar pattern of responses to the mid-PGa, i.e., increased neural responses for LC and HC conditions as compared with NC condition (p=0.002 and p=0.001, respectively) and no significant difference between LC and HC conditions (p>0.999). Interestingly, all the different AG regions showed some sensitivity to the presence of contextual information. However, whilst the ventral PGa region was positively activated (compared to rest) for NC conditions, the mid-PGp and dorsal PGa exhibited negligible activation for the NC conditions (one sample t-tests revealed ps>0.05 for NC conditions).

###### Semantic control network ROIs

We tested whether neural responses for each task-condition (NC, LC and HC) were statistically different across the different ROIs of the semantic control network. Hence, we conducted a repeated-measures ANOVA with “Region” (see below) and “Condition” (NC, LC and HC) as within-subjects factors. A significant interaction between “Region” and “Condition” revealed a different profile of engagement of the semantic control regions during the different task conditions [F(12,252)=2.48, p=0.004, *η*p^2^=0.106]. The effect of shift of semantic context was predominantly observed in the right hemisphere. The right IFG (∼BA47) [LC>NC (p=0.004), HC vs. NC (p>0.999), and LC>HC (p=0.017)] and the right dAG [LC>NC (p=0.004), HC vs. NC (p>0.999), and LC>HC (p=0.007)] showed increased responses for LC>HC conditions.

With the exception of the left ACC/pre-SMA (∼BA32/8/6) [LC>NC (p=0.017), HC vs. NC (p>0.999), and LC>HC (p=0.015)], the effect of semantic update was not observed in the left hemisphere [left IFG (∼BA45/44): LC>NC (p=0.047), HC vs. NC (p=0.145), LC vs. HC (p=0.324); left pMTG (∼BA21/37/20): LC vs. NC (p=0.917), HC vs. NC (p=0.230), LC vs. HC (p=0.99); left IFG (∼BA47): LC>NC (p=0.043), HC vs. NC (p=0.289), and LC vs. HC (p=0.573)]. Finally, the right IFG (∼BA44/45) showed a context integration effect [LC>NC (p=0.01), and HC>NC (p=0.088), and LC vs. HC (p=0.102)].

The independent ROI analyses revealed three key findings: first, the gyral distribution of semantic task activation in the temporal lobe supported previous research and revealed novel insights on the functional specialisation of the ATL for time-extended combinatorial processes. As expected, we observed a bilateral involvement of the ATL in semantic processing (Hoffman et al., 2015; Humphreys et al., 2015; Rice et al., 2015; Visser and Lambon Ralph, 2011). Interestingly, the effect of semantic coherence (HC>LC) was observed in the MTG. The aSTG and ITG showed a general effect of semantic integration (HC&LC>NC), as they were engaged more strongly – or uniquely in the case of ITG – for those condition preceded by a context paragraph (i.e., LC and HC conditions). Secondly, the re-setting of the semantic system (LC>HC) engages brain structures generally recruited when semantic processing requires increased executive control demands (Noonan et al., 2013). However, differently from previous studies, these effects are mainly observed in the right hemisphere. Finally, as found in many previous studies (Humphreys et al., 2019; Humphreys and Lambon Ralph, 2015, 2017; Seghier et al., 2010), the response profile in AG was found to shift rapidly and quickly across the AG region. The anterior ventral portion (vPGa) was sensitive to the semantic coherence of the information to be integrated, whereas instead dPGa and mid-PGp were not. Instead, these two AG sub-regions showed a semantic integration effect as they were engaged only for those condition preceded by a semantic contextual support (i.e., LC and HC conditions).

##### Whole-brain univariate analyses of the differences between experimental conditions

###### Semantic Integration effect (or formation of the conceptual gestalt)

Both HC and LC conditions generated very similar, overlapping neural semantic networks including different portions of the dorsal ATL, extending to posterior portions of the superior temporal sulcus (pSTS), ventral portions of the AG (vPGa) and bilateral IFG (Figure 3A and Table S3). Furthermore, the overlap was also observed in correspondence of “extra-semantic” areas, i.e., areas that are not referred in the literature to semantic cognition in particular, such as the left hippocampus, the right AG, medial superior frontal gyrus (mSFG) and the precuneus/posterior cingulate cortex (PCC).

**Figure 3.**
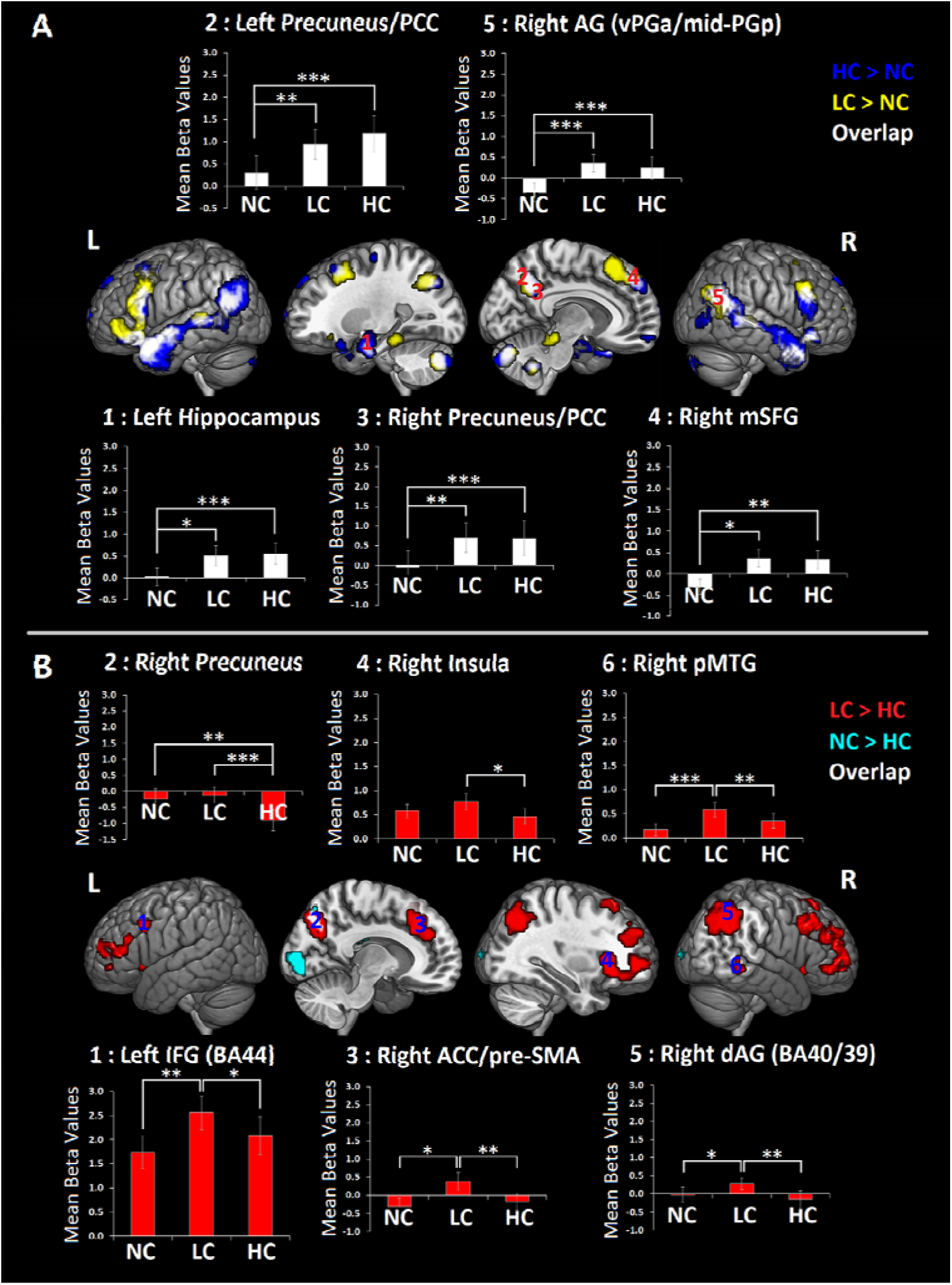
GLM results for **(A)** the semantic integration effect (HC&LC>NC) and five ROIs (derived from the peaks of activation in areas of overlap). GLM results for **(B)** the shift of semantic and task context effects (LC>HC and NC>HC, respectively) and six ROIs derived from the peaks of activation. Error bars correspond to SE.

Given neural responses within the semantic network were extensively investigated (see Figure 2), the following ROI analysis focussed on overlapping extra-semantic regions in order to reveal their role in semantic integration. Therefore, we conducted a repeated-measures ANOVA with “Region” [left hippocampus (coordinate: x=−24, y=−9, z=−24), the right AG (∼vPGa/mid-PGp; coordinate: x=54, y=−60, z=27), the right medial superior frontal gyrus (mSFG) (∼BA9/32; coordinate: x=6, y=48, z=39) and the precuneus/posterior cingulate cortex (PCC) (coordinate: x=±9, y=−54, z=36)] and “Condition” (NC, LC and HC) as within-subjects factors. The “Region” and “Condition” interaction was not significant, suggesting a similar profile of engagement of these regions during the different task conditions [F(8,168)=1.337, p=0.229, *η*p^2^=0.06]. A significant effect of “Condition” [F(2,42)=22.003, p<0.001, *η*p^2^=0.512], revealed increased neural responses in these areas for LC and HC conditions as compared with NC condition (ps<0.001). However, no significant difference was observed between HC and LC conditions (p>0.999). Interestingly, as for two portions of the AG (Figure 2B), the semantic integration effect in these areas was driven by qualitative rather than quantitative differences between conditions. In fact, these brain areas were all positively engaged (compared to rest) for LC and HC conditions (see Figure 3A), but not for NC condition (one sample t-tests revealed all ps>0.05 for NC condition). To summarise these results, integration of meaning across a narrative engages areas of the semantic network (Lambon Ralph et al., 2017) as well as other brain regions. Unlike the semantic network regions, these extra-semantic areas are recruited only when contextual information can be integrated.

###### Shift of semantic context (or update of the conceptual gestalt) and shift of task context

The ACC/pre-SMA and the precuneus were activated more for the LC than HC condition. These regions were joined by other frontoparietal MD regions (Duncan, 2010) (e.g., lateral IFG, superior parietal areas and insula) and the right pMTG (Figure 3B and Table S3). We conducted a repeated-measures ANOVA with “Region” [dorsal right precuneus (∼BA7; coordinate: x=12, y=−72, z=42), right dAG (∼BA40/39; coordinate: x=42, y=−51, z=45), right ACC/pre-SMA (∼BA32/8; coordinate: x=3, y=30, z=42), left IFG (∼BA44; x=−45, y=12, z=33), right insula (∼BA47; coordinate: x=30, y=24, z=6), right pMTG (∼BA21/20; coordinate: x=60, y=−42, z=−6)] and “Condition” (NC, LC and HC) as within-subjects factors. A significant interaction between “Region” and “Condition” revealed a different profile of engagement of the selected ROIs during the different task conditions [F(10,210)=8.504, p<0.001, *η*p^2^=0.288]. Bonferroni-corrected post-hoc comparisons revealed that neural responses in the left IFG (∼BA44) were increased for LC conditions as compared to HC and NC conditions (p<0.036 and p<0.003, respectively), but no significant difference was observed between the HC and NC conditions (p=0.164). A similar profile of differential engagement was observed in right pMTG, right dAG and right ACC/pre-SMA [right pMTG: LC>NC (p<0.001), HC vs. NC (p=0.125), and LC>HC (p=0.003); right dAG: LC>NC (p=0.011), HC vs. NC (p=0.652), and LC>HC (p=0.006); right ACC/pre-SMA: LC>NC (p=0.016), HC vs. NC (p>0.999), and LC>HC (p=0.007)]. Activity in a dorsal portions of the right precuneus (∼BA7) was increased for LC and NC conditions as compared to HC conditions (p<0.001 and p=0.002, respectively). However, there was no significant difference between LC and NC conditions (p>0.999). Finally, neural activity in the right insula (∼BA47) was increased for LC conditions as compared to HC conditions (p=0.049), whilst no significant differences were observed between NC conditions as compared to LC and HC (p=0.243 and p=0.366, respectively).

Interestingly, in contrast with what we observed for the left IFG, right insula and right pMTG, the LC>HC differential activations measured in the dorsal right precuneus, right dAG and ACC/preSMA were all due to differential negligible or task-negative activation patterns (one sample t-tests revealed all ps>0.05 for NC condition). The HC>LC contrast did not reveal significant results.

Finally, we directly compared the NC>HC condition to establish which brain regions are important for task switches into language from a non-language task (number reading) and therefore to identify possible similarities between neuro-computations supporting shifts of semantic and task contexts. This contrast activated a right lateralised set of higher-order visual regions, including also a portion of the right precuneus activated by the LC>HC contrast (see Figure 3B and Table S3). To summarise, resetting the conceptual gestalt elicits a robust activation of a bilateral set of frontal regions and the right pMTG. The right precuneus was less deactivated for LC and NC conditions as compared with HC condition, a pattern that mirrored that of RTs (see Figure 1).

#### 3.2.2. Task group spatial ICA results

##### Task-related FNs

ICA was used to explore which semantic and extra-semantic areas, revealed by univariate analyses, exhibited yoked activations – i.e., constituted FNs rather than independent areas. ICA identified 22 ICs, of which 5 exhibited significant sensitivity to our task conditions: These were a semantic/language network (SLN), an executive control network (ECN) including fronto-parietal regions, a higher visual network (HVN), a primary visual network (PVN), and a DMN (Figure 4 and Table S4).

**Figure 4.**
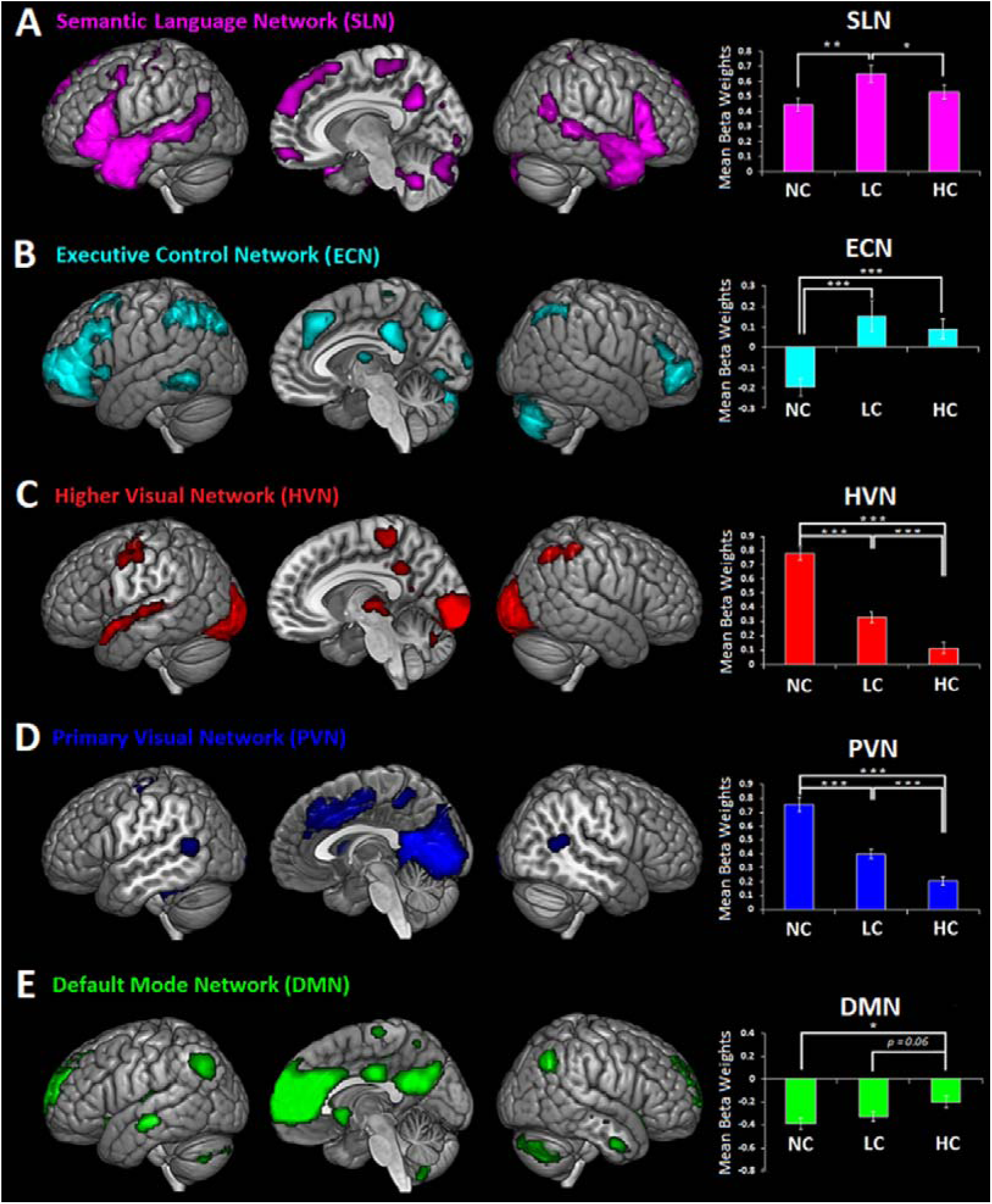
Task-related FNs and beta weights’ results. **(A)** Semantic language network (SLN); **(B)** Executive control network (ECN); **(C)** Higher visual network (HVN); **(D)** Primary visual network (PVN); **(E)** Default-mode network (DMN). Error bars correspond to SE.

As well as a bilateral set of semantic brain areas, the SLN included extra-semantic regions (e.g., hippocampus), suggesting that these regions are recruited together with semantic areas to support time-extended semantic cognition. We conducted a repeated-measures ANOVA on the beta weight values. The results revealed a significant interaction between “Network” (SLN, ECN, HVN and DMN) and “Condition” (NC, LC and HC) within-subjects factors [F(8,168)=42.698, p<0.001, *η*p^2^=0.670], suggesting different engagement of the networks in the different task conditions. In fact, the SLN was positively engaged by all three conditions (in comparison to rest), but that it was most active when the semantic context shifted (i.e., LC condition) [LC>NC (p=0.007), LC>HC (p=0.032)]. A related pattern was observed in the ECN which was positively engaged by both the HC and LC conditions, i.e., when a previous semantic context was available for integration [LC>NC (p<0.001), HC>NC (p<0.001)], though this was independent of whether the context was semantically congruent with the target or not [HC vs. LC (p=0.774)]. The two visual networks (HVN and PVN), containing occipital but also attentional control regions, were recruited most heavily for the NC condition, in which there was a change in the cognitive task [HVN: NC>LC (p<0.001), NC>HC (p<0.001) and LC>HC (p<0.001); PVN: NC>LC (p<0.001), NC>HC (p<0.001) and LC>HC (p=0.001)].

Finally, unlike the four other components, the DMN was deactivated with respect to rest. It exhibited sensitivity to the task conditions, in that it was least deactivated in the HC condition [HC>NC (p=0.025), HC>LC (p=0.063), and NC vs. LC (p=0.88)]. This pattern of deactivation mirrors the task performance (see Figure 1), in which RTs for the target narrative were slowest for the LC and NC condition. Thus this result might reflect the common pattern that the degree of deactivation in the DMN is often correlated with task/item difficulty and other measures of behavioural stability (Esterman et al., 2013; Humphreys et al., 2015; Kucyi et al., 2016; Vatansever et al., 2017b). A negative correlation between averaged DMN TCs (averaged across runs and time-courses) and averaged RTs was observed in our study, without however reaching statistical significance (*r*=−0.099, p=0.662).

##### FNC analysis

Given our interest in studying the interaction between the SLN and other networks involved in time-extended semantic cognition, we computed a FNC analysis (see Figure 5). Significant positive correlations were observed between the TCs of DMN and SLN (*r*=0.13) and between DMN and ECN (*r*=0.23). Instead, the DMN was negatively correlated with HVN (*r*=−0.22). The ECN, the network not engaged during changes of task context, but only during semantic integration, showed a significant negative correlation with both HVN and PVN (*r*=−0.17 and *r*=−0.18, respectively), i.e., the networks maximally engaged during changes of task context (NC condition). In contrast with ECN, the SLN was positively correlated with both the HVN and PVN (*r*=0.28 and *r*=0.21, respectively). Finally, the TCs of the two visual networks were positively correlated (*r*=0.41).

**Figure 5.**
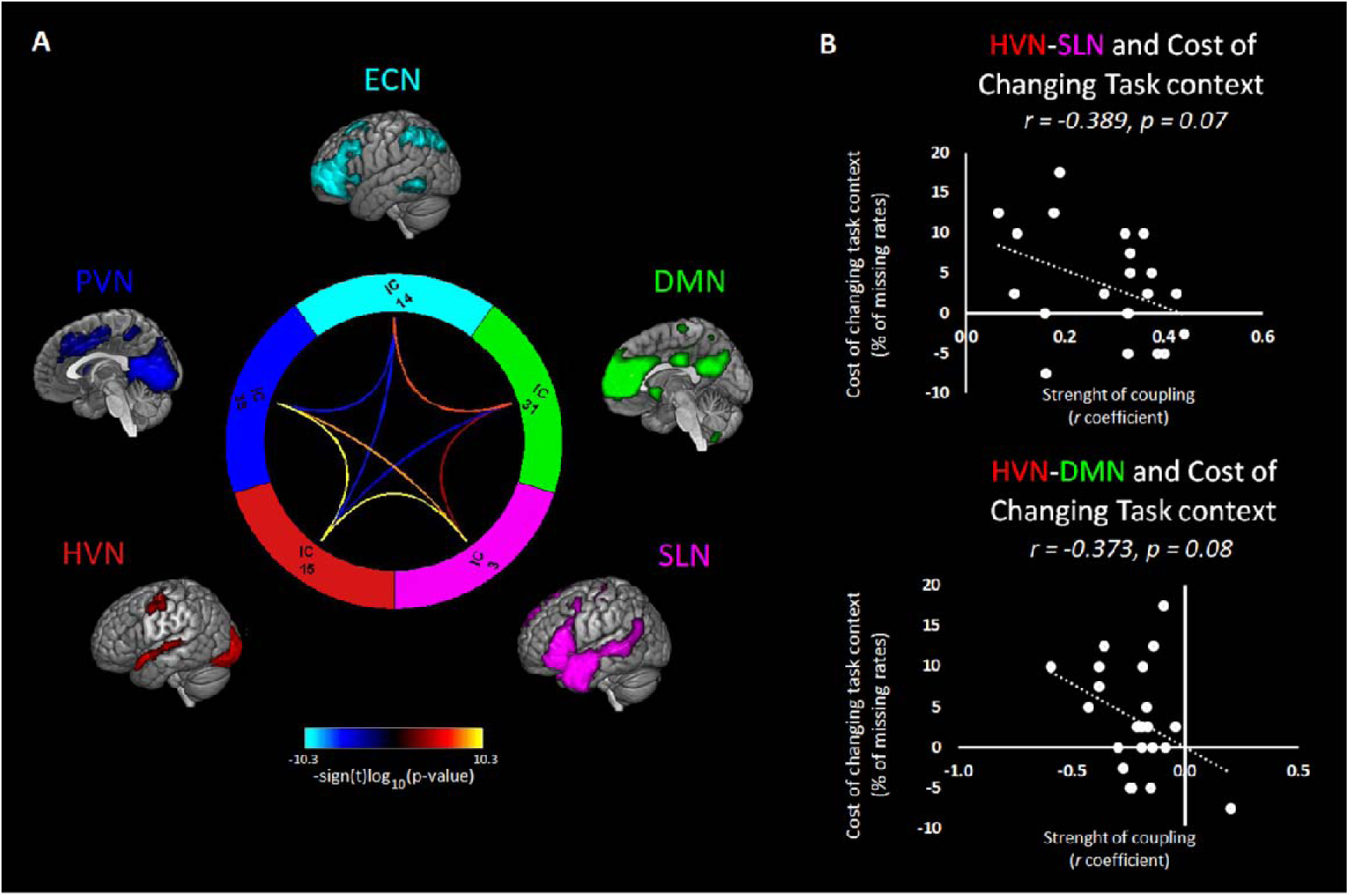
FNC results. **(A)** Connectogram of the FNC results: Significant pairwise correlations (p<0.05) were corrected for multiple comparisons (FDR, α□=□0.01). Significance and direction of each pairwise correlation is displayed as the −sign(t)log10(p); and **(B)** Correlations between strength of coupling and the behavioural cost associated with changes of task context.

These results may suggest that time-extended semantic integration is supported by both the SLN and ECN, interacting with the visual networks in different ways. Unlike the SLN, ECN recruitment is anticorrelated with the visual FNs, being strongly recruited when switching from number to text reading (NC conditions, i.e., shift of task context). Thus, it might be that switching efficiently from a non-semantic to a semantic task requires the activation of semantic and language areas (SLN) and, at the same time, the disengagement of brain regions involved in integration of contextual information (ECN). To test this hypothesis, we assessed whether the HVN-SLN positive coupling and the HVN-ECN negative coupling were negatively correlated with the behavioral cost of changing task context (operationalized as the difference between NC and HC conditions calculated for both % of missing rates and RTs). Given recent findings (Crittenden et al., 2015; Smith et al., 2018), we also tested whether this correlation could be observed with the HVN-DMN coupling.

Results revealed that the stronger the positive coupling between HVN and SLN, the smaller the behavioural cost associated with changes of task context (NC>HC % of missing rates) (*r*=−0.389, p=0.07) (Figure 5 B). Despite the correlation between the HVN-ECN coupling and the same behavioural cost was not significant (*r*=−0.121, p=0.59), we found that the HVN-DMN negative coupling was negatively correlated with the behavioural cost associated with changes of task context (NC>HC % of missing rates) (*r*=−0.373, p=0.08). The same correlation analyses were conducted with behavioural costs measured as RTs. However, these analyses did not reveal significant results or trends toward significance.

## 4. Discussion

Using both univariate and multivariate data-driven (ICA) approaches we revealed the brain regions and the networks supporting time-extended formation and updating of conceptual representations. Unlike prior investigations (Hasson et al., 2008; Lerner et al., 2011; Simony et al., 2016; Tylen et al., 2015), in this fMRI study we established which neural networks are primarily evoked by the formation and updating of semantic representations and we distinguished them from those that support other non-semantic processes required during narrative reading. The main findings on the functional specialisation of brain areas within and outside the classical semantic network are discussed below, followed by a discussion on the FNs, i.e., how semantic and extra-semantic regions (i.e., regions outside the classical semantic network) are recruited together during meaning formation and update.

### The Semantic Network

We hypothesised that the building of the conceptual gestalt would be supported by two interactive neural systems, reflecting representational and control aspects of semantic cognition, respectively. In accord with our hypothesis and previous findings (Hasson et al., 2008; Lerner et al., 2011; Silbert et al., 2014), our results revealed that integrating semantic information during narrative processing engages a bilateral set of frontal, parietal and temporal brain structures, known as the “semantic network” (Lambon Ralph et al., 2017). Within this network we identified hubs for formation of coherent concepts and control regions for context-sensitive regulation of semantic information.

### Hubs for the formation of time-extended conceptual gestalt: the role of ATL

A first important result obtained from the univariate analysis (whole-brain and independent ROI analyses) is that the ATL supports time-extended combinatorial processes in addition to basic semantic combinations shown in previous studies (Brennan and Pylkkanen, 2012; Visser and Lambon Ralph, 2011), corroborating the hypothesis that this region is a key hub for the formation of conceptual representations (Noppeney and Price, 2004; Patterson et al., 2007; Lambon Ralph et al., 2017; Vandenberghe et al., 2002). In particular, the effect of semantic coherence (HC>LC) was observed in the MTG, suggesting that this subregion of the ATL may have a crucial role in the formation of time-extended conceptual gestalt in verbal tasks. Future studies will have to assess whether this effect is observed in other subregions of the ATL when meaningful stimuli are presented in non-verbal modalities. In contrast, the aSTG and the ITG showed a general effect of semantic integration (HC&LC>NC) as they were engaged more strongly – or uniquely in the case of ITG – for those condition preceded by a context paragraph (i.e., LC and HC conditions).

The graded differential distribution of activation patterns observed within the ATL may reflect differences of structural connectivity between ATL subregions and primary input/output areas (Binney et al., 2012). The lack of sensitivity for NC conditions in the ITG indicates that neural activity in this ATL subregion is modulated by the presence of contextual information. This effect may be determined because of direct connections (via entorhinal cortex) with regions supporting the formation of contextual memories, such as in the parahippocampal gyrus and the hippocampus (Bar et al., 2008; Davachi, 2006; Dundon et al., 2018). A recent study has demonstrated the interplay between semantic processing in ATL and information encoding/retrieval in the hippocampus during the formation/retrieval of contextual memories (Griffiths et al., 2019). Accordingly, it is possible that these anatomical connections allow these regions to cooperate for the formation and retrieval of semantic and contextual information conveyed in the narrative (e.g., episodic details) (Ranganath and Ritchey, 2012), facilitating the flow of neocortical information into the hippocampus during encoding and the propagation of hippocampal retrieval signals into the ATL during retrieval. Future experimental studies combining high-temporal and high-spatial resolution techniques will be needed to test this possibility.

### Brain regions for the update of time-extended conceptual gestalt: the contribution of the right hemisphere

A second important result is that increased semantic integration demands induced by the need to update of the semantic gestalt elicit bilateral activation of ventral and dorsal portions of the frontal lobe (Figure 3B). This result is in accord with previous studies that have employed tasks requiring multi-item combinations (Silbert et al., 2014; Tylen et al., 2015). Conversely, it differs from others that have utilised semantic association tasks, where the involvement of the IFG was mostly left lateralised (Humphreys et al., 2015; Noonan et al., 2013).

The involvement of the right hemisphere during natural-like language processing has been attributed to the complexity of the input. In other words, as language input gets increasingly complex, there is increasing involvement of right hemisphere homologues to classic left hemisphere language areas (Jung-Beeman, 2005). Particularly, the right hemisphere activation may become prominent and sustained when words and sentences are presented in a narrative context, and may here reflect coherence and inference at the propositional level and beyond, when readers make connections between sentences and paragraphs to form a coherent conceptual gestalt. This interpretation is in accord with the view that the right hemisphere, as compared to the left, would be involved in processing global aspects of linguistic contents (Hickok and Poeppel, 2007; Poeppel, 2003; St George et al., 1999).

Nevertheless, this proposal does not entirely fit with our results since univariate results revealed that some key nodes within the semantic control network (i.e., dAG, pMTG and ventral IFG) show enhanced responses in the right, but not in the left hemisphere when integration requires major revision of the semantic context (LC>HC contrast). Hence, despite the modulation of control regions in the right hemisphere is in accord with the findings in the literature (Jung-Beeman, 2005; Xu et al., 2005), it is not clear what might have determined a shift in the lateralisation of the effect related to the update of the semantic gestalt. One possibility is that this effect is not observed in the left hemisphere because neural responses in these regions may have been too transient to be captured by fMRI.

Consistent with previous findings, the involvement of the right parietal and frontal regions may reflect the recruitment of domain-general WM (Gajardo-Vidal et al., 2018; Vigneau et al., 2011) and inhibitory control mechanisms (Aron et al., 2004) applied when incongruences are detected across paragraphs of text. Particularly, the right ventral IFG (BA47) and the right insula might enact the sustained suppression of the incorrect interpretation of the ambiguous word and all the words semantically related to it, after a change of semantic context is detected (Mason and Just, 2007).

The role of the left pMTG in semantic tasks has been related to the modulation of semantic activation to focus on aspects of meaning that are appropriate to the task or context (Noonan et al., 2013). Accordingly, we were expecting that this region would also respond to shifts of semantic context (i.e., during the update the semantic gestalt). However, this effect was observed in the right pMTG that has different structural and functional connectivity properties as compared to the left pMTG (Vassal et al., 2016; Xu et al., 2015). For instance, Neurosynth (http://old.neurosynth.org/; Yarkoni et al., 2011) shows that resting-state co-activation maps from the left and right pMTG regions have different patterns of connectivity with the ATL. That is, the left, but not the right pMTG, shows intrinsic connectivity with the ATL semantic hub. Instead, the right pMTG shows connectivity with regions that, except for the left pMTG, constitute a right lateralised fronto-parietal network. This observation along with evidence that this area is not involved in semantic tasks (Noonan et al., 2013), unless they require formation of time-extended contextual associations (Xu et al., 2005), may indicate that the right pMTG plays a critical role in capturing changes of context-sensitive meaning over longer periods of time, possibly by integrating information across WM (frontal and parietal corticies) and semantic networks (left pMTG).

### The role of the left AG: semantic hub or buffering system?

Our results reveal that a portion of the left AG (vPGa) supports meaning formation during time-extended semantic cognition similar to the MTG. In fact, an effect of semantic coherence was observed in the anterior ventral portion of the left AG (vPGa). This result aligns with the proposal that the left AG has a crucial role for the formation and retrieval of semantic representations (Binder and Desai, 2011; Binder et al., 2009; Geschwind, 1972). However, this general semantic role for the AG does not fit with a series of findings from neurological patients that have reported semantic impairments after ATL but not parietal damage (Lambon Ralph et al., 2017) and the demonstrations of equal (de) activation for non-semantic and semantic tasks in this region (Humphreys et al., 2015; Humphreys et al., 2019; Humphreys and Lambon Ralph, 2015, 2017).

Recent compelling evidence has led to an alternative proposal on the role of the left AG that would reconcile neuroimaging and neuropsychological findings. Rather than a hub for semantic integration, the left AG might support a domain-general mechanism that buffers time, context-, and space-varying inputs (Humphreys et al., 2019; Humphreys and Lambon Ralph, 2017). Since buffering becomes relevant only when the task requires combinations across multiple (internal or external) items (e.g., during encoding, integration, recollection, etc.), it is not surprising that left ventral AG is positively engaged in our study and in other semantic tasks that involve integration of information across multiple items (Price and Mechelli, 2005; van der Linden et al., 2017).

### Beyond the Semantic Network

Besides brain areas implicated in semantic cognition, we also expected to observe additional brain regions and networks reflecting the demands posed by forming and updating conceptual contexts. In accord with previous findings (Hasson et al., 2008; Lerner et al., 2011; Price, 2012; Silbert et al., 2014; Simony et al., 2016; Vigneau et al., 2006; Xu et al., 2005), a set of brain regions comprising the hippocampus, the precuneus/PCC and the DMN AG region (i.e., mid-PGp) responded to semantic integration (LC&HC>NC contrast). Similar to the mid-PGp, these other regions have been identified as nodes of both task-positive (e.g., episodic memory, mind-wandering, etc.) and task-negative (DMN) networks, and have been related to multiple cognitive functions, including semantic cognition (Krieger-Redwood et al., 2016). In accord with what previous studies have found when contrasting intact (similar to our HC condition) versus scrambled (similar to NC condition) narratives (Hasson et al., 2008; Lerner et al., 2011; Simony et al., 2016), we revealed that activity in the hippocampus, the precuneus/PCC and the mid-PGp (see above) was modulated by context integration. However, contrary to what it has been suggested, activity in these areas was not modulated by the semantic coherence per se, but rather by the presence of the contextual support. Without saying that these are fully independent neural processes, the intact vs. scrambled narrative contrast, used in many previous investigations, does not necessarily distinguish between brain areas crucial for the formation of a semantic gestalt (meaning integration) and those important for scene construction processes, i.e., those processes that allow linking incoming cues about the current context (e.g., time, space cues) into a schema representation or situation model (for a discussion on these two different systems see also (Ranganath and Ritchey, 2012). Our study allowed us to draw out this distinction for the first time. Specifically, we compared brain activity elicited by coherent narratives (HC condition) not only against conditions without a contextual support (i.e., without a situation model, NC condition), but also against conditions that, despite having a contextual support, nevertheless required an update of the ongoing semantic representation (i.e., the meaning associated to words and their combinations, see LC condition).

Therefore, we propose that whilst activity in the ATL is likely to reflect semantic integration per se (see above), activity in DMN regions (i.e., hippocampus, the precuneus/PCC and the mid-PGp) may support processes needed to construct associations between the information conveyed in the target and context paragraphs (Davachi, 2006; Maguire et al., 1999; Ranganath and Ritchey, 2012; Spaniol et al., 2009; Staudigl and Hanslmayr, 2013).

Finally, contrary to what has been suggested by previous investigations (Crittenden et al., 2015; Smith et al., 2018), DMN regions (e.g., precuneus/PCC and mSFG) were not increasingly engaged by shifts of semantic or task contexts. These inconsistencies might be due to substantial differences in the tasks employed in our and other studies. Being a naturalistic-like task, the reading task was quite “passive” and resetting the task context did not require the cognitive manipulation (retrieval, inhibition, etc.) of instructions/rules associated to the task to be performed. Instead, switching between the highly novel tasks employed in previous studies (Crittenden et al., 2015; Smith et al., 2018) necessarily requires this sort of process, given that each stimulus domain (i.e., stimuli depicting people, buildings, words) was associated to two possible classification rules (male/female and old/young for face stimuli; skyscraper/cottage and inside/outside view for building stimuli; first letter and last letter for word stimuli). Hence, the task-switch activity observed in DMN regions in previous studies may reflect the retrieval of task rules, rather than reinstatement and assessment of contextual representations. A related possibility is that given the novelty of these tasks (learned prior to scanning), the DMN region activations reflect episodic retrieval of the task instructions.

### The Functional Networks

ICA was employed in addition to univariate analysis to reveal how different brain regions were recruited for semantic integration. In the present study, ICA revealed a network sensitive to semantic control demands (SLN), a network sensitive to context integration (ECN) and two networks sensitive to domain-general attentional control demands (HVN and PVN). Finally, ICA revealed also a DMN network modulated by task-condition difficulty.

### SLN and ECN: semantic control and working memory processes

In accord to previous research work (Geranmayeh et al., 2014; Price, 2012; Silbert et al., 2014; Simony et al., 2016; Vigneau et al., 2006), and an influential model of semantic cognition (Lambon Ralph et al., 2017), we expected two task-positive networks to support semantic processing and executive control. We were expecting these two networks to be modulated by semantic integration (LC&HC>NC) and shifts of semantic context (LC>HC). According to our predictions, ICA revealed a SLN and ECN, both positively engaged during the semantic task conditions.

The SLN network, including semantic, but also MD and other extra-semantic areas (e.g., hippocampus), was maximally recruited during shifts of semantic context (update of the semantic gestalt). This finding suggests that updating the information in the semantic system requires the orchestration of different neurocomputations possibly including WM, semantic processing and domain-general executive control.

The ECN network, including a set of fronto-parietal and medial regions, was also positively engaged during the reading task. However, rather than being sensitive to variations of semantic context (LC>HC), the ECN was sensitive to context integration in general (LC&HC>NC). That is, this network was positively recruited when the target conditions could be integrated with a previous context, independently of whether the context was highly congruent with the target. Instead, this network was disengaged when information could not be integrated with a previous context. The spatial distribution of this network - including some key nodes identified as WM areas from GLM analysis - and its sensitivity to contextual integration is consistent with a WM function (Vatansever et al., 2017a).

### The visual networks support domain-general attentional control mechanisms

ICA analysis revealed that two additional networks were positively engaged during the reading task. These networks, including visual areas, but also other cortical and subcortical control regions (e.g., IFG, pMTG, precentral gyrus, putamen, etc.), were strongly engaged during changes of task and semantic context. This result is particularly interesting because it shows that sensory-dorsal and posterior attentional networks are involved in narrative reading and support control functions that are important, but not specific, for semantic integration.

Furthermore, FNC analyses revealed that HVN and PVN were both positively connected to the SLN and negatively connected to the DMN and ECN. We proposed that this coupling might reflect the need of strongly engaging semantic areas after the number reading task, and at the same time, disengaging areas not relevant for the switch of task context, e.g., ECN. Our result revealed that switching efficiently between tasks was related to stronger negative coupling between HVN and DMN and at the same time, enhanced positive coupling between HVN and SLN. According to the DMN-downregulation hypothesis described above, the negative correlations between task-positive HVN and task-negative DMN (Fox et al., 2005; Vincent et al., 2008) may suggest that the visual network was constantly engaged during the task (except during rest). This is not surprising since HVN and PVN include visual areas and other control structures that are constantly active during the reading tasks and maximally engaged during increased control demands.

### The role of DMN during narrative processing

We expected recruitment of a DMN including hippocampus, AG (mid-PGp), precuneus/PCC, and other medial prefrontal structures (Andrews-Hanna et al., 2010). According to the hypothesis that DMN supports semantic integration (Binder et al., 2009), we should have observed task-positive responses or some sensitiveness to semantic manipulations in the task. However, like previous findings, the DMN showed the typical task-negative response (Geranmayeh et al., 2014; Humphreys and Lambon Ralph, 2015; Wirth et al., 2011). Whilst we acknowledge that the DMN involvement or absence in semantic processing cannot be decided on the basis of this evidence alone, the observation that DMN activity was not modulated by semantic integration is hard to reconcile with the semantic hypothesis (Binder et al., 2009). Furthermore, replicating previous studies (Humphreys et al., 2015), we observed similar DMN task-negative responses during semantic and non-semantic processing alike (see Figure S1C). This finding is clearly at odds with the hypothesis that the primary function of the DMN is representing and integrating semantic information.

A second hypothesis is that the DMN supports episodic retrieval and buffering (Rugg and Vilberg, 2013; van der Linden et al., 2017; Vatansever et al., 2017a; Vilberg and Rugg, 2008). If so, then one would expect to observe an activation profile similar to the ECN: positively engaged only for conditions allowing integration (LC and HC). Conversely, not only was the DMN negatively engaged during meaning integration (HC and LC), but most importantly, it did not show differential responses between conditions preceded (LC) and not preceded by a contextual support (NC). Thus, this result is inconsistent with the proposal that DMN supports episodic retrieval and buffering during narrative reading.

A final and third hypothesis is that the DMN is involved in cognitive transitions by reinstating context-relevant information (Crittenden et al., 2015; Smith et al., 2018). As such one would expect the DMN to be sensitive to a major switch to a new task (NC condition), when a completely different context representation is reawakened. In contrast to this prediction, we found larger task-negative activations for NC as compared with HC.

It is worth noting that the evidence that we provide here might seem contradictory with previous reports suggesting a key DMN role in narrative processing (Ames et al., 2015; Baldassano et al., 2017; Baldassano et al., 2018; Chen et al., 2017a; Lerner et al., 2011; Simony et al., 2016; Yeshurun et al., 2017). One possibility is that we might have failed to observe DMN responses reflecting semantic integration in the current study because the stimuli employed were shorter than the much longer narratives used in previous studies (Baldassano et al., 2017; Baldassano et al., 2018; Chen et al., 2017a; Lerner et al., 2011; Yeshurun et al., 2017). There is, however, evidence suggesting that the involvement of DMN areas may not relate to the temporal duration of the stimuli, but rather to the presence of informational context. In fact, stimuli with the same informational content but different durations, recruit the DMN to the same extent (Baldassano et al., 2017). Furthermore, short narratives with an interleaved design (similar to the one used here) recruit DMN regions, such as PCC and medial prefrontal cortex, responding to context integration (Ames et al., 2015).

Our GLM results revealed a positive engagement of DMN regions during context integration (i.e., HC&LC>NC). This set of regions included the same regions that have been found in other studies (Hasson et al., 2008; Simony et al., 2016). In apparent contrast with the GLM results, however, our ICA results revealed that the engagement of the DMN, including the regions noted above, reflected a task-negative engagement sensitive to task difficulty. An explanation for these apparently contrasting results might reside in a common mechanism between “cognitive ease” and “narrative processing”. For instance, one could hypothesise that during easier semantic integration (HC condition) as compared to harder semantic integration (LC) the DMN might support the formation of a schema representation. This type of cognitive processes, typically used for task-negative internal mentation (e.g., autobiographical memory and future planning), would facilitate the building of an internal model of the narrative and therefore facilitate its comprehension (i.e., the formation of a semantic gestalt). Therefore, despite the mean activity in DMN regions co-varies with an overall attentional state (and/or internal vs. external mentation) - in accord with the pattern of RT responses - yet at the same time the local neural circuits may be processing information related to the stimuli (formation of schema representation and conceptual gestalt). This hypothesis, however, contrasts with the observation that no significant difference between conditions with (LC condition) and without context-related schema representations (NC condition) was observed.

An alternative explanation of why some nodes of the DMN (e.g., ATL, hippocampus) were positively engaged (compared to rest) during our task conditions (GLM results), whilst the DMN overall activity is task-negative and co-varies with task difficulty, is that the DMN is not homogeneous and fractionates depending on the task contrasts (Axelrod et al., 2017; Buckner et al., 2008; Buckner and DiNicola, 2019; Humphreys et al., 2015). In parallel with previous investigations, we found that the same regions (ATL, mid-PGp and hippocampus) were aligned with the SLN during narrative reading but became a part of the extended DMN when semantic integration is not required (e.g., during number reading see Figure S1A-B). Finally, the observation that DMN responses mirror RTs (Figure 1) (see also Humphreys and Lambon Ralph, 2017; Vatansever et al., 2017a) accords with the proposal that, when not critical for the task at hand, some brain regions may be deactivated, proportional to task difficulty, to save metabolic energy whilst preserving performance (Attwell and Laughlin, 2001).

## ACKNOWLEDGEMENTS

This work was supported by a Medical Research Council Programme Grant (MR/R023883/1), an European Research Council Advanced Grant (GAP: 670428 - BRAIN2MIND_NEUROCOMP) and a Postdoctoral Fellowship from the European Union’s Horizon 2020 research and innovation programme, under the Marie Sklodowska-Curie grant agreement No 658341.

