## Supplementary material for "Revealing the neural networks that extract conceptual gestalts from continuously evolving or changing semantic contexts": SI

Francesca M. Branzi or Matthew A. Lambon Ralph

**This Supplementary file includes:**

Figure S1

Tables S1 to S4

Captions for Tables S1 to S4

References for SI reference citations

**Figure S1.** GLM results were FDR−corrected voxel−wise at a statistical threshold of q < 0.05, and a contiguity threshold ≥ 30 voxels. Task > Rest (blue) and Rest > Task (green) activations in (A) non-semantic (number context condition) and in (B) semantic (narrative context condition) reading tasks. (C) Mean beta weights’ results relative to the DMN for the semantic and non-semantic reading tasks (context conditions, i.e., SR and NSR respectively). The engagement of the DMN did not differ in SR and NSR conditions [t(21)=0.387, p=0.703].

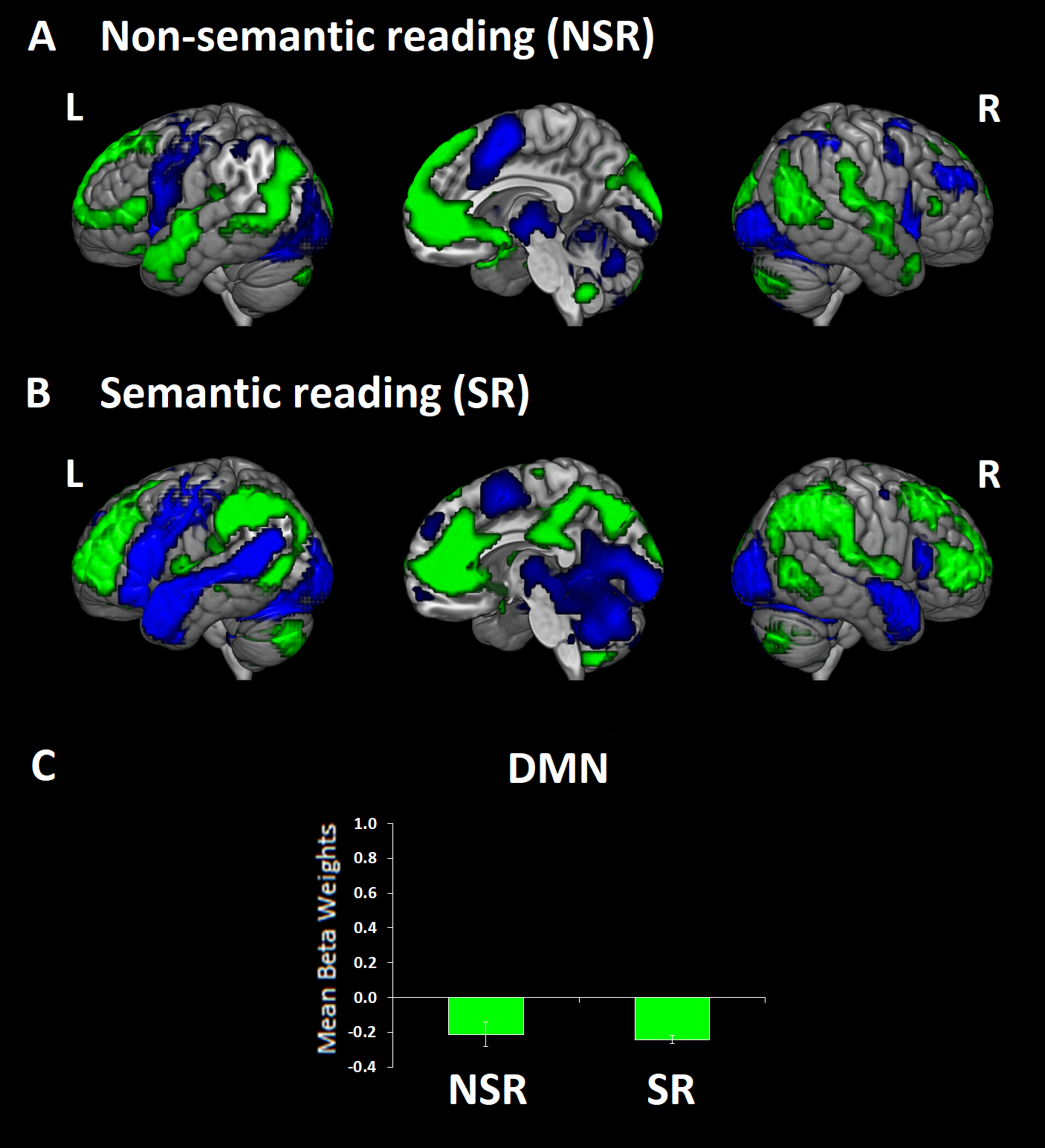

**Table S1. S**timuli for Low−Congruent (LC) and High−Congruent (HC) conditions (Context and Target paragraphs). The shift of semantic context after Low−congruent contexts was expected to be perceived after the critical ambiguous word, here depicted in red.

| **Narrative code** | **LC Context** | **HC Context** | **Target** |
| --- | --- | --- | --- |
| 1 | Paul is quite talkative, but this evening he was completely engaged in checking his new suit. When he noticed that the hem of the trousers was unstitched, he realised he had to fix it before going to the charity event. | Tom is quite chatty, but this morning he was totally absorbed in listening to the tennis match. When one of the players broke the other’s serve, he realised his favourite player would win soon. | ***While he was thinking about how excited he was, the speaker on the radio said that the ball hit the ground on the line. His favourite player had won the match and so he went to celebrate.*** |
| 2 | Gina visited Rome. During the first few days, she wanted to explore the city. After a long walk along the river, she found a place to sit and get some rest. This seat is not very comfortable, she thought, but it’s better than nothing. | Amy visited London. During the first few hours, she wanted to explore the city. In the centre, she found an ATM to withdraw some money. She wanted to buy something. The exchange rate is not very good, she thought, but it’s better than nothing. | ***While she was there the bank phoned to tell her that her account was blocked again. She had to wait one day before buying anything. She could not believe it had happened again.*** |
| 3 | Since she was young she had always hated baseball. This is because her cousin used to talk about this sport all the time. This is why she never wanted to go to the stadium, until she met her boyfriend. | Since she was a child she had always hated vampires and nocturnal animals. This is because her brother used to talk about these disgusting creatures all the time. This is why she never wanted to walk in the dark, until she met her boyfriend. | ***Her boyfriend was popular in her city because he used to have a bat living in his house. I remember I used to see it flying around the neighbourhood at night. In fact, these animals do not like the light.*** |
| 4 | This famous driver was explaining that his car had a problem which made it slow. It seemed that it was a fuel system problem. The team had to work hard to fix it. Jamie was watching the TV very carefully. | This famous psychologist was explaining the causes of wrongful convictions throughout history. It seemed that it was a social minority problem. He had to work hard to prove this. Joe was watching the TV very carefully. | ***The expert said that race is one aspect of social biases that leads to discriminatory behaviours. This psychologist was saying that these biases can affect people’s lives dramatically.*** |
| 5 | After being a little unsure, John finally went to the hairdresser to cut his long hair. When he came out with the new cut, his hair was short, but he was satisfied. | After being a little doubtful, Luke finally went to the music shop to buy something new. When he came out with the new guitar, his wallet was empty, but he knew that it was for a good reason. | ***He still needed the band to accept him and he hoped that now he might be invited to take part in the next concert.*** |
| 6 | The two boxers were in front of each other, ready to fight. We were watching attentively. We knew about the rules. Before the match started, the speaker took the microphone and made an announcement. | Jessica and Rob were in front of each other, ready to dance. We were watching attentively. We knew about their engagement. When the music started, he took the microphone and made an announcement. | ***At that moment the camera captured the ring. It was shiny and had a big diamond. They would get married for sure. We thought we would only see this kind of thing in the movies.*** |
| 7 | We discovered many things that week. We were told about the rainforest and the environmental disaster in Peru. Indeed, three major oil spills had drastically affected the trees of the rainforest. | We learnt many things that day. We learnt about elephants and the natural disaster in Tanzania. Indeed, two major climatic changes had drastically affected the number of elephants. | ***We also learnt that trunks can be used by elephants to rub an itchy eye. Moreover, these animals use their trunk to threaten and to throw objects, and as snorkels when swimming in water.*** |
| 8 | The tour was interesting. We were told that the station was built during an expedition led by an explorer on his arrival at the beginning of the past century. Henceforth, every year there is a ceremony during which a new flag is raised in a different place. | The tour was interesting. We were told that the station was erected during an expedition led by an explorer on his arrival at the beginning of the past century. Henceforth, every year there is a ceremony during which the ice’s drift is measured. | ***What we found particularly interesting about this is that the pole is moving at a rate of roughly ten metres per year.*** |
| 9 | The garden is a great place to stay in spring and she spends many hours there. She takes care of the plants and she places traps near the small holes in the ground. Unfortunately, she knows very well that it is necessary. | The garden is a nice place to stay in summer and she spends some time there. She takes care of the flowers and she covers her skin. Unfortunately, she knows very well that it is necessary. | ***Given the large number of moles on her skin she has to be careful with the sun. The doctor told her that she should wear a large hat on her head while she is doing these activities.*** |
| 10 | On the first day of school he had to go in front of the class to introduce himself. The professor knew that this experience was quite challenging for him. However, it could not be avoided. | On the second day of treatment she had to apply a lotion to the patient’s eyes. She knew that this lotion was quite irritating. However, it could not be avoided. | ***She knew that the pupils dilate when someone is about to cry. She learnt it during the first year of medical training. She was feeling sorry for him and she did not know what to do.*** |
| 11 | Emma cultivates many different types of flowers and since last month she has worked hard on the garden outside her house. It required a lot of work, but we all agree that it was worth doing. | Julie makes many different types of lamps and since last year she has worked hard on the lighting outside her house. She invited me to come over to see her work. | ***When I went to visit her, the bulbs outside the house were not working at all. When I went back home it was dark and I had to use the light from my mobile phone to find my way.*** |
| 12 | Today I saw Becky and she told me she was coming back from her first Pilates class. She told me that she wanted to do some exercise to tone up her body. She is generally quite lazy, so I was quite surprised. | Yesterday I saw Grace and she told me that she was going to sort out her clothes that day. She told me that she wanted to do something to improve the appearance of her room. She is generally quite lazy, so I was quite surprised. | ***I saw that her chest of drawers was full of neatly folded clothes. I realised that she was really committed this time. At that moment, I thought I could follow her example.*** |
| 13 | Lucy was listening to the news on the radio when she recognised her colleague’s voice. She was glad to hear his interview about the research project on Alzheimer's disease. He received many questions as his view contrasted with other experts’ views. | Ruth was reading the news on her computer when she recognised her own words. She was glad to read her interview about the research project on Alzheimer's disease. While she was reading, she noticed that Outlook was flickering. | ***The account was full of emails. Her colleagues from Europe were sending their best wishes and asking about the details of the new project. Some of them wanted to collaborate.*** |
| 14 | When I met her yesterday, we started to talk about the fact that after years of political discussion, Parliament had finally voted to establish a new law to ban the sale of electronic cigarettes in the United Kingdom. | I met her yesterday in the theatre because her seat was next to mine. We started to talk about the fact that after ten years of incessant performing, she had decided to leave for a while in order to work on a new character. | ***We stopped talking since we realized that the next act was going to be a very talented young actor. We concentrated on the theatrical performance.*** |
| 15 | We were watching the players on the green. He was explaining to me that the goal of the game is to roll biased balls so that they stop close to a smaller ball. I was listening to him carefully because I really wanted to learn. | We were preparing the lunch in the kitchen. He was explaining to me that the secret of preparing a good salad is to have a dressing made with balsamic vinegar. I was listening to him carefully because I really wanted to learn. | ***Look at that bowl, you almost made it fall and it is full of salad and oil! I was paying too much attention to what he was saying and I almost spilled the oil over the floor.*** |
| 16 | Laura has announced that she is going to launch her own business. She plans to hire someone with expertise in material sciences as well as in global markets. It will take some time for her to have everything sorted out. | My best friend Julie has decided that she is going to come back to Manchester. She plans to buy a flat in the same neighbourhood where I live. It will take some time for her to have everything sorted out, but I am really glad about her decision. | ***Her company has always been a pleasure for me for different reasons. She has always been a great and supportive friend.*** |
| 17 | I went with Mum to the beauty salon because I should do something for my hands. I received a treatment that improved my hands’ appearance. This included a coat of polish. When we went back home, I sat in the lounge and then I started to look at my hands. | I went with Mum to the shopping centre because I should do something for the walls in my house. We bought a hammer and a screwdriver for hanging up pictures. When we went back home, we went to the lounge and then we started to use my purchases. | ***Also Dad was there and at some point he noticed that the nails were not hammered fully into the wall; hence I tried to help Dad by holding the nail in position while he was hammering it into the wall again.*** |
| 18 | I have been there many times before, but we took a nice cruise on the sea last year. We stayed on the boat for two weeks visiting the islands around the coast. | I have been there many times before, but we took a superb wine tour last year. We spent two weeks in the countryside, visiting vineyards around the area. | ***Furthermore, the port wine was to die for. We did not know this wine before. It is incredibly sweet and rich in taste and it pairs wonderfully with desserts.*** |
| 19 | I recently saw his girlfriend. I have to say he has been very lucky this time. I tried very hard, but I struggled not to stare at her. She is very amusing and intelligent. | I finally saw his girlfriend's new present. I have to say she has been very lucky this time. I tried very hard, but I struggled not to stare at it. It is a very beautiful bracelet. | ***She had a charm bracelet that is similar to the one I bought for my sister. It is a double silver chain that has small gold and silver pendants with little shining diamonds.*** |
| 20 | Yesterday I attempted to make sweet bread with figs and other dry fruits. In Egypt I discovered my passion for figs. I love to eat them especially when fresh, but I also like to cook them. | I finally looked at the calendar and I attempted to make a plan for this summer. In Spain I discovered my passion for travelling. I love to do that especially in the summer, but I also like to travel during winter. | ***Today, after several attempts, I finally made a date for the next trip. As well as my friends, I would like to ask to my sister and her boyfriend to come with us.*** |
| 21 | I remember this guy from school. He was a peculiar one. He had some vision problems, in the sense that he was short-sighted. | I remember this guy from school. He was a peculiar one. He had some problems with wine, in the sense that he could hardly drink anything. | ***Once he had some glasses of wine. At the end he was so drunk that we had to carry him home. I remember his Dad and his mother were mad. They probably thought it was our fault.*** |
| 22 | My little niece convinced us to go to the circus last weekend. It’s not that I didn’t want to, but I was not interested in seeing the acrobats. However, we went along with my parents, and in the end I was glad they convinced me to go. | My little nephew convinced us to go to the shopping centre last Sunday. It’s not that I didn’t want to, but I was not interested in doing any clothes shopping. However, we went along with my parents, and in the end I was glad they convinced me to go. | ***While we were there, I realised that the jumper that I was wearing had a hole. It was cold and I needed to get a new one. I was really upset because I bought it just a few days ago.*** |
| 23 | I want to introduce my friend Claire to you next week. You have many things in common. For instance, she is a terrific chef. She is an expert in cooking different types of food and especially liver. | I want to introduce my sister Eve to you next week. You have many things in common. For instance, she is a terrific musician. She is an expert in playing different types of instruments, especially the piano. | ***Moreover, one of the things she likes most is organ music. She recently went to various organ recitals and I am keen to go with her one of these days.*** |
| 24 | We were told to come over to see their new plants and flowers behind the house. However, we were not really impressed about them. | We were invited to come over to see the size of their new boat. However, we were not really impressed about it. | ***In terms of yards, it is twenty yards in length. So it is not small, but we think that they could have organized the space better.*** |
| 25 | My father was born before the end of the war. They used to live in the countryside when he was two years old. They were living outside the city because of the high risk of bombings. | My father was born many years ago. This picture was taken in the countryside when he was two years old. He wanted to be held by my grandmother all the time and he liked to play with her neck. | ***The arms of my father were round my grandmother’s neck in this picture. He was hugging her. In this picture you can also see my grandfather. He is the tallest man standing close to them.*** |
| 26 | I recently went there with my friends. It was such a wonderful place. We stayed there for two weeks. We also visited a castle with rustic wooden walls and roofs. I liked everything about that place. | I recently went there with my friends. It was such a beautiful place. We stayed two weeks in a hotel. In our room there was plenty of sunlight. I liked everything about the room. | ***If I have to choose a particular thing, I really liked the beams of light that came through the big skylight of our room and the way that they scattered around us.*** |
| 27 | Recently, Maya was recognised as a national talent. However, we realized earlier that our daughter had a scientific mind. By the age of four, she could do very complicated mathematical calculations. | Last week, Rachel was recognised as a national talent. Nevertheless, we realized earlier that our sister was brilliant. By the age of three, she became famous because of her incredibly fast hands. | ***By only using the digits of one of her hands she was able to solve the Rubik's Cube in a few minutes.*** |
| 28 | She has been busy recently, because she is studying hard for an exam in macroeconomics. She told to me that it refers to a branch of economics dealing with the economy as a whole, rather than individual businesses. | She has been quite busy recently, because she is studying very hard for an exam. Since I haven’t seen her lately, I decided to invite her to stay for dinner and I checked the food in the fridge. | ***Then I realised that the market was going to close soon. I had to go out quickly to buy something for dinner or I would not have anything tasty to prepare.*** |
| 29 | It has not been easy getting together to play cards recently, but Jane finally invited us to come over this weekend. She has a beautiful house where we will play different card games. | It has been complicated getting together for vacations recently, but Elsa finally invited us to come over this weekend. She has a beautiful house with a great view on the bay where we will get some rest. | ***Honestly, nothing could be better than the bridge’s view from her beautiful terrace. I am sure that we will have a great time in San Francisco.*** |
| 30 | This morning I prepared breakfast for my wife. I turned the radio on and I decided to make pancakes. I started to mix egg flour and milk in a bowl, stirring until it was smooth. | Yesterday morning I woke up before my wife. I turned the radio on and I heard a famous cricket player being interviewed by a radio station because of his recent achievements. | ***I was surprised because the batter has made a half century of runs in one hundred straight games yesterday. The radio was saying that it's been nearly a year since his streak began.*** |
| 31 | She had guests for lunch and she was entertaining them with her recent adventures in Costa Rica. She said that she did a lot of bird watching and she learnt a lot about those animals. However, she thought sometimes it was quite difficult. | Mary had guests for dinner and she was entertaining them with her culinary skills. She said that she had attended cooking classes and had learnt how to cook different cuts of meat. However, she thought sometimes it was quite difficult. | ***“You need to pay attention. It’s rare when its colour is dark red with some juice flowing.” Her husband was trying to explain to her how to cook a rare steak. She was not good at it.*** |
| 32 | He met his colleague at noon. He had a working lunch with him to examine their collection of photos. He saw a powerful woman with a married actor. He wanted to break the scandal to the world. | He met his future wife at noon. They had the sample lunch to go over the rehearsal menu and the desserts. They still had to decide on the best desserts. They wanted to find the best ice-cream. | ***He thought that he would have wanted another scoop of ice-cream, but he didn’t dare to ask the waiter.*** |
| 33 | In the lodge there was just the sound of chirping bush animals. It lasted for six hours. The noise was so loud that we could not hear each other. At the beginning I could not really fall asleep. After a few days I got used to it. | In our house there was just the sound of the TV broadcasting the match. The match was so long that we could not watch any other programme. At the beginning I could not really understand it. After some time I got used to it. | ***My husband did not care at all. In fact, he was absorbed in watching the cricket match on the TV, which is the thing he likes best in the world.*** |
| 34 | My cousin owns a property in Italy. He often visits in spring. The last time we went there, one of his kids hurt his leg whilst playing football. When he started to cry loudly, everyone got really scared. | My brother owns a property in France where he farms different animals. The last year we went there one of his cows became ill. When it started to cry loudly, everyone got really scared. | ***You should have seen the calf. I felt sorry for it. It was so scared that it quickly went to hide behind the cow's belly.*** |
| 35 | When I arrived there with my brother, we immediately went to explore the island. The weather was sunny and very warm, so we decided to go to the beach. Coconuts were no longer hanging in the trees, so we decided to sit under them. | When I arrived there with my brother, we immediately went to explore the vineyard. The weather was nice and warm, so we started to harvest grapes. After one hour it turned out to be quite hot. My hands were aching, so we decided to stop. | ***While we were resting there, my brother noticed that the palm of my hand was bleeding. I probably touched something sharp and I hadn’t realised.*** |
| 36 | During the weekend Tom loves to sit and watch motorbike races. That night the TV news announced that his favourite racer was going to win the competition. However, another rider moved ahead and remained in the lead until the end. | During the evening Noel loves to sit on the sofa and watch the TV with his dog. That night the TV news announced that the movie was going to start soon. Hence, at that moment, he bent his knees, he took the blanket and waited for his dog. | ***While he was there, he noticed that his lap was not comfortable enough for his dog. It did not want to stay on it. Since the dog preferred to lie on the sofa, he stretched out his legs.*** |
| 37 | Hopefully your aunt will not be offended, but I think that we will not be able to arrive in time. It snowed a lot yesterday and now there is a lot of traffic. Even if I try to find a solution, I am sure we are not going to get out of this problem. | I hope that she will not be upset, but I think we will not be able to bring our strawberry dessert. I was in a rush yesterday and I forgot to buy the ice cream. Even if I try to find a solution, I am sure we are not going to get out of this problem. | ***I was very upset. It got worse when I realised that the jam that I prepared with the strawberries from our garden spilled over the backseats. We were not going to have any dessert today.*** |
| 38 | We started to talk about the global outcry over the escape of the gorilla from the cage. The cage was not strong enough. Hence, the gorilla was able to damage it and escape. | We ordered some drinks and we started to talk about the outcry over the proposal to prohibit selling alcohol. The main problem is that many people are not able to limit their drinking. | ***The bar was full of people that day. We were having some drinks and at one point the barman offered us a beer because we are frequent customers.*** |
| 39 | I went with Lucy to the stadium to watch our favourite rugby team. The opposing team scored many points despite the fact they were missing two players. As you could imagine, shortly after we arrived I got quite nervous. | Despite it being a rainy day, I went with Sara to the park last week. We went to the tobacco shop because I wanted to buy cigarettes but it was closed. As you could imagine, shortly after we arrived I got quite stressed. | ***Moreover, I realised that the matches were completely wet because of the rain and I could not light my cigarette. I asked her if she had a lighter but she told me she quit smoking.*** |
| 40 | We got lost in the countryside. I stopped the car and asked where I could find the way back. A pedestrian told me that we had to go straight and then turn to the right. Indeed, on the left side there was a blind turn. Then we returned to the car and we started to drive. | We went to buy some cutlery. I parked the car and asked where I could find some knives. An employee told me that they were at the end of the corridor. We took them and then we went to pay. Then we returned to the car and we started to drive. | ***We were driving for some time and I realised that the forks that we bought were not on the backseat as we thought. We forgot to put them in the car.*** |

**Table S2.** MNI coordinates for ROIs derived from the literature.

| **ROI groups** | **Study** | **Type of Analysis** | **x** | **y** | **z** | **Location** |
| --- | --- | --- | --- | --- | --- | --- |
| Temporal lobe | Rice et al., 2015 | Activation likelihood clusters from the GingerALE analyses (Verbal input modality studies) | ±48 | 16 | −28 | aSTG (BA38) |
|  |  |  | ±58 | 8 | −20 | MTG (BA21) |
|  |  |  | ±36 | 12 | −34 | ITG (BA20) |
| Left AG | Humphreys et al., 2019 | Task ICA | −48 | −69 | 36 | L mid−PGp |
|  |  |  | −33 | −72 | 46 | L dPGa |
|  |  |  | −51 | −54 | 21 | L vPGa |
| Semantic control network | Noonan et al., 2013 | Activation likelihood clusters from the GingerALE analyses (High > Low Semantic Control Studies) | −45 | 19 | 21 | L IFG (BA45/44) |
|  |  |  | −35 | 22 | −11 | L IFG (BA47) |
|  |  |  | −55 | −50 | −5 | L pMTG (BA21/37/20) |
|  |  |  | −3 | 16 | 49 | L ACC/pre−SMA (BA32/ 24/8/6) |
|  |  |  | 36 | −63 | 39 | R dAG (BA39/7) |
|  |  |  | 35 | 22 | −11 | R IFG (BA47) |
|  |  |  | 47 | 22 | 29 | R IFG (BA44/45) |

L=left; R=right; a=anterior, p=posterior; d=dorsal; v=ventral; STG=superior temporal gyrus; MTG=middle temporal gyrus; ITG=inferior temporal gyrus; AG=angular gyrus; IFG=inferior frontal gyrus; ACC/pre−SMA=anterior cingulate cortex/pre−supplementary motor area.

**Table S3.** MNI coordinates and locations of the activation peaks from the GLM analyses. GLM results were FDR−corrected voxel−wise at a statistical threshold of *q*<0.05, and a contiguity threshold ≥30 voxels.

| **Contrast** | **cluster size** | **T** | **x** | **y** | **z** | **Location** |
| --- | --- | --- | --- | --- | --- | --- |
|  | 17923 | 23.86 | 12 | −87 | −3 | R Lingual Gyrus |
| Semantic Targets > Number Context |  |  |  |  |  |  |
|  |  | 23.49 | -6 | -93 | -6 | L Calcarine Gyrus |
|  |  | 19.52 | 18 | -93 | 6 | R Superior Occipital Gyrus |
|  | 167 | 7.61 | -3 | 66 | -12 | L Mid Orbital Gyrus |
|  |  | 3.77 | -6 | 48 | -18 | L Rectal Gyrus |
|  | 472 | 5.61 | -9 | 54 | 42 | L Superior Frontal Gyrus |
|  |  | 5.26 | -6 | 18 | 63 | L Posterior-Medial Frontal |
|  |  | 5.15 | -6 | 57 | 33 | L Superior Medial Gyrus |
|  | 310 | 4.34 | -45 | -21 | 21 | L Rolandic Operculum |
|  |  | 3.72 | -39 | -6 | 18 | L Rolandic Operculum |
|  | 30 | 3.27 | -3 | -9 | 51 | L Posterior-Medial Frontal |
|  | 20217 | 19.12 | −18 | −90 | −9 | L Occipital cortex (Area hOc3v [V3v]) |
| Semantic Targets > rest |  |  |  |  |  |  |
|  |  | 16.9 | 18 | −93 | −6 | R Calcarine Gyrus |
|  |  | 16.27 | −6 | −93 | −6 | L Calcarine Gyrus |
|  | 61 | 6.41 | −3 | 66 | −15 | L Mid Orbital Gyrus |
|  | 115 | 4.28 | 33 | 0 | 54 | R Middle Frontal Gyrus |
|  |  | 3.38 | 45 | 3 | 57 | R Middle Frontal Gyrus |
|  | 52 | 4.11 | -9 | 51 | 36 | L Superior Medial Gyrus |
|  | 1302 | 8.93 | 48 | 9 | −30 | R Medial Temporal Pole |
| High-Congruent > No-Context |  |  |  |  |  |  |
|  |  | 8.13 | 51 | 6 | −21 | R Medial Temporal Pole |
|  |  | 8.12 | 66 | −6 | −18 | R Middle Temporal Gyrus |
|  | 397 | 8.33 | −9 | −51 | 36 | L Precuneus |
|  |  | 6.58 | 9 | −57 | 36 | R Precuneus |
|  | 2354 | 8.25 | −48 | −66 | 27 | L Angular Gyrus (PGp) |
|  |  | 7.93 | −54 | −60 | 24 | L Angular Gyrus (PGa) |
|  |  | 7.26 | −39 | −66 | 36 | L Angular Gyrus (PGp) |
|  | 394 | 7.83 | 21 | −81 | −42 | R Cerebelum |
|  | 142 | 6.27 | 60 | 21 | 27 | R IFG p. Opercularis |
|  | 193 | 5.64 | −21 | −9 | −21 | L Hippocampus |
|  |  | 4.57 | −36 | −12 | −30 | L Fusiform Gyrus |
|  | 228 | 5.22 | −21 | −78 | −39 | L Cerebelum |
|  | 82 | 5.03 | 6 | −57 | −45 | R Cerebelum |
|  |  | 4.05 | −6 | −60 | −45 | L Cerebelum |
|  | 345 | 4.92 | 3 | 48 | 39 | L Superior Medial Gyrus |
|  |  | 4.05 | 15 | 54 | 42 | R Superior Medial Gyrus |
|  |  | 3.91 | −9 | 21 | 57 | L Posterior-Medial Frontal |
|  | 46 | 4.71 | −9 | 69 | 6 | L Superior Medial Gyrus |
|  | 217 | 4.64 | −33 | 21 | 51 | L Middle Frontal Gyrus |
|  |  | 4.13 | −36 | 6 | 48 | L Precentral Gyrus |
|  | 45 | 4.51 | 0 | 60 | −15 | L Rectal Gyrus |
|  |  | 3.41 | 18 | 63 | −12 | R Superior Orbital Gyrus |
|  | 155 | 4.43 | −54 | 21 | 12 | L IFG p. Triangularis |
|  |  | 4.13 | −54 | 21 | 27 | L IFG p. Triangularis |
|  | 38 | 4.08 | 24 | −9 | −21 | R Hippocampus |
|  | 31 | 4 | 51 | −78 | 3 | R Middle Occipital Gyrus |
|  |  | 3.46 | 54 | −72 | 9 | R Middle Temporal Gyrus |
|  | 49 | 3.88 | 33 | −33 | −18 | R Fusiform Gyrus |
|  |  | 3.61 | 39 | −27 | −18 | R Fusiform Gyrus |
|  | 162 | 3.88 | −27 | −15 | 63 | L Precentral Gyrus |
|  |  | 3.59 | −57 | −18 | 36 | L Postcentral Gyrus |
|  |  | 3.36 | −54 | −18 | 48 | L Postcentral Gyrus |
|  | 76 | 3.76 | −39 | −24 | 18 | L Rolandic Operculum |
|  | 292 | 7.02 | 15 | −78 | −33 | R Cerebelum |
| Low-Congruent > No-Context |  |  |  |  |  |  |
|  | 863 | 6.92 | 39 | −60 | 30 | R Angular Gyrus (PGa) |
|  |  | 4.93 | 54 | −60 | 27 | R Angular Gyrus (PGp) |
|  |  | 4.52 | 54 | −36 | 3 | R Middle Temporal Gyrus |
|  | 322 | 6.21 | 3 | 24 | 57 | L Posterior-Medial Frontal |
|  |  | 3.8 | 6 | 45 | 42 | R Superior Medial Gyrus |
|  | 252 | 5.91 | 48 | 12 | −30 | R Medial Temporal Pole |
|  |  | 5.06 | 57 | 6 | −30 | R Middle Temporal Gyrus |
|  |  | 4.25 | 48 | 27 | −15 | R IFG p. Orbitalis |
|  | 154 | 5.32 | 57 | 21 | 33 | R IFG p. Opercularis |
|  | 203 | 5.27 | −39 | −24 | 18 | L Rolandic Operculum |
|  |  | 5.07 | −33 | −33 | 18 | L Rolandic Operculum |
|  |  | 2.91 | −54 | −18 | 6 | L Superior Temporal Gyrus |
|  | 336 | 5.24 | −6 | −57 | −42 | L Cerebelum |
|  |  | 4.96 | −12 | −75 | −30 | L Cerebelum |
|  |  | 4.92 | 9 | −54 | −45 | R Cerebelum |
|  | 810 | 5.03 | −27 | −69 | 42 | L Inferior Parietal Lobule |
|  |  | 4.99 | −60 | −60 | 21 | L Angular Gyrus (PGp) |
|  |  | 4.77 | −33 | −69 | 33 | L Middle Occipital Gyrus |
|  | 1402 | 4.85 | −45 | 21 | −24 | L Temporal Pole |
|  |  | 4.7 | −27 | 12 | 48 | L Middle Frontal Gyrus |
|  |  | 4.65 | −60 | −6 | −9 | L Middle Temporal Gyrus |
|  | 300 | 4.56 | 12 | −54 | 33 | R Precuneus |
|  |  | 4.25 | −3 | −60 | 45 | L Precuneus |
|  |  | 4.2 | 6 | −60 | 42 | R Precuneus |
|  | 110 | 4.17 | −12 | −33 | −15 | L Cerebelum |
|  |  | 3.81 | −24 | −36 | −18 | L Fusiform Gyrus |
|  |  | 3.18 | −12 | −45 | −24 | L Cerebelum |
|  | 66 | 4.01 | −39 | −12 | −27 | L Inferior Temporal Gyrus |
|  |  | 3.5 | −30 | −9 | −27 | L ParaHippocampal Gyrus |
|  | 70 | 3.97 | 9 | −30 | −15 | R Cerebelum |
|  |  | 3.39 | 18 | −36 | −15 | R Cerebelum |
|  |  | 2.69 | 27 | −33 | −18 | R Fusiform Gyrus |
|  | 49 | 3.19 | 33 | 12 | 45 | R Middle Frontal Gyrus |
|  | 30 | 3.16 | 33 | 57 | 12 | R Superior Frontal Gyrus |
|  | 67 | 3.15 | −39 | −21 | 39 | L Postcentral Gyrus |
|  |  | 2.77 | −54 | −18 | 36 | L Postcentral Gyrus |
|  | 430 | 6.49 | 3 | 30 | 42 | L Superior Medial Gyrus/ACC-preSMA |
| Low-Congruent > |  |  |  |  |  |  |
| High-Congruent |  |  |  |  |  |  |
|  |  | 5.54 | 9 | 27 | 36 | R Middle Cingulate Cortex |
|  |  | 5.19 | 3 | 24 | 54 | L Posterior-Medial Frontal |
|  | 1123 | 6.36 | 12 | −72 | 42 | R Precuneus |
|  |  | 5.87 | 48 | −54 | 33 | R Angular Gyrus (PGa) |
|  |  | 5.25 | 42 | −51 | 45 | R Inferior Parietal Lobule (IPS) |
|  | 76 | 5.22 | −30 | 33 | −12 | L IFG p. Orbitalis |
|  |  | 3.55 | −39 | 48 | −6 | L Middle Orbital Gyrus |
|  | 995 | 5.14 | 39 | 51 | −3 | R Middle Orbital Gyrus |
|  |  | 4.95 | 30 | 27 | −9 | R IFG p. Orbitalis |
|  |  | 4.86 | 30 | 24 | 6 | R Insula Lobe |
|  | 53 | 4.79 | 60 | −42 | −6 | R Middle Temporal Gyrus |
|  | 194 | 4.77 | −45 | 12 | 33 | L Precentral Gyrus |
|  |  | 4.35 | −42 | 3 | 27 | L Precentral Gyrus |
|  | 164 | 4.73 | −51 | 39 | 9 | L IFG p. Triangularis |
|  |  | 4.53 | −54 | 30 | 24 | L IFG p. Triangularis |
|  |  | 4.1 | −36 | 57 | 18 | L Middle Frontal Gyrus |
|  | 77 | 3.92 | −36 | 15 | −3 | L Insula Lobe |
|  |  | 3.88 | −39 | 15 | 9 | L Insula Lobe |
|  | 39 | 3.69 | 24 | 27 | 60 | R Superior Frontal Gyrus |
|  |  | 3.5 | 30 | 21 | 57 | R Middle Frontal Gyrus |
|  | 148 | 6.53 | 24 | −96 | 6 | R Middle Occipital Gyrus |
| No-Context > High-Congruent |  |  |  |  |  |  |
|  |  | 5.99 | 15 | −87 | −6 | R Lingual Gyrus |
|  | 50 | 6.05 | 15 | −66 | 36 | R Precuneus |
|  |  | 4.35 | 9 | −72 | 54 | R Precuneus/Superior Parietal Lobe |
|  | 55 | 5.63 | 0 | −30 | 24 | Posterior Cingulate Cortex |

L=left; R=right; IFG=inferior frontal gyrus

**Table S4.** MNI coordinates and locations of the activation peaks for each task−related FN. The results were FWE−corrected voxel−wise at a statistical threshold of *p*<0.05, and a contiguity threshold ≥30 voxels.

| **Component** | **cluster size** | **T** | **x** | **y** | **z** | **Location** |
| --- | --- | --- | --- | --- | --- | --- |
| **SLN** | 5601 | 34.49 | -51 | -9 | -12 | L Superior Temporal Gyrus |
|  |  | 32.23 | -57 | -6 | -21 | L Middle Temporal Gyrus |
|  |  | 28.71 | -51 | 24 | 12 | L IFG (p. Triangularis) |
|  | 443 | 22.11 | 6 | -57 | 30 | R Precuneus |
|  |  | 16.28 | -6 | -57 | 33 | L Precuneus |
|  |  | 9.95 | -18 | -48 | 36 | L Precuneus |
|  | 1142 | 22.09 | -3 | 54 | 27 | L Superior Medial Gyrus |
|  |  | 21.86 | -9 | 51 | 36 | L Superior Medial Gyrus |
|  |  | 19.83 | -6 | 15 | 63 | L Posterior-Medial Frontal |
|  | 2657 | 21.09 | 48 | 33 | -15 | R IFG (p. Orbitalis) |
|  |  | 20.42 | 45 | 15 | -24 | R Temporal Pole |
|  |  | 19.61 | 57 | 24 | -3 | R IFG (p. Orbitalis) |
|  | 232 | 20.28 | 6 | -57 | -42 | R Cerebelum (IX) |
|  |  | 18.43 | -6 | -57 | -45 | L Cerebelum (IX) |
|  |  | 9.53 | 12 | -42 | -33 | R Cerebelum |
|  | 579 | 19.76 | 21 | -75 | -30 | R Cerebelum (Crus 1) |
|  |  | 17.98 | 21 | -81 | -39 | R Cerebelum (Crus 2) |
|  |  | 14.17 | 18 | -81 | -21 | R Cerebelum (Crus 1) |
|  | 1017 | 15.66 | -12 | -27 | 66 | L Paracentral Lobule |
|  |  | 14.16 | -33 | -21 | 66 | L Precentral Gyrus |
|  |  | 13.33 | -33 | -36 | 63 | L Postcentral Gyrus |
|  | 155 | 13.17 | 6 | 57 | -15 | R Rectal Gyrus |
|  | 194 | 12.81 | -18 | -75 | -30 | L Cerebelum (Crus 1) |
|  |  | 12.48 | -18 | -81 | -39 | L Cerebelum (Crus 2) |
|  | 60 | 11.82 | 15 | -84 | 9 | R Calcarine Gyrus |
|  |  | 9.75 | 12 | -72 | 12 | R Calcarine Gyrus |
|  | 71 | 11.25 | 33 | -93 | 15 | R Middle Occipital Gyrus |
|  |  | 10.57 | 27 | -84 | 24 | R Superior Occipital Gyrus |
|  | 53 | 10.13 | 15 | -54 | -3 | R Lingual Gyrus |
|  | 63 | 9.46 | -12 | -51 | -6 | L Lingual Gyrus |
|  | 48 | 8.37 | -39 | -48 | -18 | L Inferior Temporal Gyrus |
| **ECN** | 4313 | 30.45 | -45 | 48 | -3 | L Middle Orbital Gyrus |
|  |  | 25.96 | -33 | 54 | 0 | L Middle Frontal Gyrus |
|  |  | 25.42 | -45 | 48 | -12 | L Middle Orbital Gyrus |
|  | 1229 | 28.17 | -51 | -45 | 48 | L Inferior Parietal Lobule (IPS) |
|  |  | 24.67 | -33 | -72 | 45 | L Inferior Parietal Lobule (dPGa) |
|  |  | 23.08 | -36 | -57 | 39 | L Angular Gyrus (IPS) |
|  | 341 | 22.14 | -6 | -30 | 36 | L Middle Cingulate Cortex |
|  |  | 10.75 | -15 | -51 | 18 | L Precuneus |
|  | 1054 | 20.96 | 30 | -66 | -36 | R Cerebelum (Crus 1) |
|  |  | 19.99 | 12 | -81 | -27 | R Cerebelum (Crus 1) |
|  |  | 19.97 | 27 | -81 | -51 | R Cerebelum (Crus 2) |
|  | 263 | 16.53 | -60 | -51 | -6 | L Middle Temporal Gyrus |
|  |  | 10.03 | -63 | -33 | -15 | L Middle Temporal Gyrus |
|  | 136 | 16.38 | -6 | -72 | 42 | L Precuneus |
|  | 184 | 13.84 | 42 | -60 | 54 | R Angular Gyrus (IPS/PGa) |
|  |  | 11.78 | 51 | -48 | 54 | R Inferior Parietal Lobule (PFm) |
|  | 106 | 13.09 | -15 | -27 | 60 | Primary Motor Cortex |
|  | 593 | 12.58 | 48 | 51 | 0 | R Middle Frontal Gyrus |
|  |  | 12.25 | 45 | 48 | -12 | R IFG (p. Orbitalis) |
|  |  | 11.85 | 39 | 42 | -9 | R Middle Orbital Gyrus |
|  | 82 | 11.83 | -9 | -42 | 0 | L Lingual Gyrus |
|  | 193 | 10.38 | -9 | -102 | 12 | L Superior Occipital Gyrus |
|  |  | 9.91 | -15 | -72 | -3 | L Lingual Gyrus |
|  |  | 9.59 | -6 | -75 | -6 | L Lingual Gyrus |
|  | 34 | 9.50 | -27 | -36 | -21 | L Fusiform Gyrus |
|  | 54 | 9.00 | 63 | -51 | -6 | R Middle Temporal Gyrus |
|  |  | 8.93 | 63 | -45 | -15 | R Inferior Temporal Gyrus |
|  | 56 | 8.43 | 21 | -99 | 15 | R Superior Occipital Gyrus |
| **HVN** | 3420 | 34.08 | 15 | -90 | -3 | R Lingual Gyrus |
|  |  | 32.40 | -9 | -90 | -3 | L Calcarine Gyrus |
|  |  | 25.54 | -12 | -93 | -18 | L Lingual Gyrus |
|  | 785 | 14.26 | -18 | -30 | 0 | L Thalamus |
|  |  | 13.26 | -60 | -30 | 6 | L Middle Temporal Gyrus |
|  |  | 12.49 | -30 | -6 | 3 | L Putamen |
|  | 197 | 12.61 | 12 | -36 | 57 | R Paracentral Lobule |
|  |  | 10.49 | -6 | -33 | 57 | L Paracentral Lobule |
|  |  | 8.63 | -9 | -27 | 66 | L Paracentral Lobule |
|  | 38 | 12.03 | 24 | -30 | 0 | R Thalamus (Temporal) |
|  | 204 | 10.86 | -51 | 0 | 42 | L Precentral Gyrus |
|  |  | 9.54 | -54 | -15 | 48 | L Postcentral Gyrus |
|  |  | 8.61 | -33 | -18 | 51 | L Precentral Gyrus |
|  | 155 | 10.85 | 51 | -60 | 51 | R Angular Gyrus (PGa) |
|  |  | 8.37 | 57 | -42 | 57 | R Inferior Parietal Lobule (PFm) |
|  |  | 7.27 | 48 | -45 | 42 | R SupraMarginal Gyrus |
|  | 101 | 10.84 | 3 | -30 | 39 | R Middle Cingulate Cortex |
|  |  | 9.91 | -3 | -42 | 33 | L Posterior Cingulate Cortex |
|  | 59 | 8.64 | 18 | -57 | 42 | R Precuneus |
|  | 66 | 8.16 | -9 | 66 | 9 | L Superior Medial Gyrus |
|  |  | 7.60 | 9 | 66 | 3 | R Superior Medial Gyrus |
|  |  | 7.50 | 0 | 63 | 0 | L Superior Medial Gyrus |
| **PVN** | 5332 | 34.97 | 0 | -87 | 0 | L Calcarine Gyrus |
|  |  | 32.36 | -6 | -72 | 6 | L Calcarine Gyrus |
|  |  | 31.92 | -18 | -57 | 12 | L Calcarine Gyrus |
|  | 775 | 14.48 | 0 | 15 | 36 | L Middle Cingulate Cortex |
|  |  | 13.69 | 0 | 0 | 45 | L Middle Cingulate Cortex |
|  |  | 13.20 | 9 | 24 | 36 | R Middle Cingulate Cortex |
|  | 120 | 13.74 | 9 | 6 | 6 | R Caudate Nucleus |
|  |  | 8.51 | 12 | -15 | 12 | R Thalamus |
|  | 154 | 12.21 | -3 | -42 | 51 | L Middle Cingulate Cortex |
|  |  | 7.49 | -3 | -63 | 54 | L Precuneus |
|  | 157 | 10.93 | -39 | -24 | 51 | L Postcentral Gyrus |
|  |  | 10.56 | -36 | -24 | 69 | L Precentral Gyrus |
|  | 36 | 10.77 | -6 | 6 | 6 | L Caudate Nucleus |
|  | 40 | 10.74 | -54 | -3 | -18 | L Middle Temporal Gyrus |
|  | 108 | 10.30 | -42 | 18 | 30 | L IFG (p. Triangularis) |
|  |  | 7.62 | -36 | 6 | 36 | L Middle Frontal Gyrus |
|  | 32 | 9.81 | 60 | 18 | -9 | R Temporal Pole |
|  |  | 8.55 | 54 | 15 | -15 | R Temporal Pole |
|  | 44 | 9.53 | 42 | 3 | 42 | R Precentral Gyrus |
|  |  | 8.41 | 36 | 12 | 48 | R Middle Frontal Gyrus |
|  |  | 6.79 | 42 | 15 | 36 | R IFG (p. Opercularis) |
|  | 34 | 8.51 | -24 | 24 | 39 | L Middle Frontal Gyrus |
| **DMN** | 4772 | 42.13 | -6 | 36 | 9 | L Anterior Cingulate Cortex |
|  |  | 39.02 | -9 | 60 | 15 | L Superior Medial Gyrus |
|  |  | 35.16 | 6 | 45 | 3 | R Anterior Cingulate Cortex |
|  | 737 | 21.28 | 3 | -51 | 27 | R Posterior Cingulate Cortex |
|  |  | 19.76 | 3 | -18 | 30 | R Middle Cingulate Cortex |
|  |  | 16.73 | -6 | -63 | 24 | L Cuneus |
|  | 101 | 16.96 | -66 | -21 | -12 | L Middle Temporal Gyrus |
|  | 85 | 15.44 | -33 | 15 | -18 | L IFG (p.Orbitalis) / Insula Lobe |
|  | 197 | 15.12 | -48 | -63 | 36 | L Angular Gyrus (Pga/PGp) |
|  | 111 | 13.02 | 54 | -60 | 39 | R Inferior Parietal Lobule |
|  | 324 | 12.99 | 30 | -81 | -30 | R Cerebelum (Crus 1) |
|  |  | 11.96 | 39 | -75 | -39 | R Cerebelum (Crus 2) |
|  |  | 11.31 | 45 | -54 | 42 | R Inferior Parietal Lobule |
|  | 90 | 12.17 | 45 | -63 | -42 | R Cerebelum (Crus 2) |
|  |  | 8.81 | -33 | -81 | -33 | L Cerebelum (Crus 2) |
|  | 71 | 11.97 | 30 | 18 | -15 | R Insula Lobe |
|  | 245 | 11.81 | -24 | -24 | -15 | L Subiculum |
|  |  | 10.99 | -51 | -12 | 0 | L Superior Temporal Gyrus |
|  |  | 9.80 | -54 | -27 | 12 | L Superior Temporal Gyrus |
|  | 54 | 11.65 | 51 | -3 | -33 | R Inferior Temporal Gyrus |
|  | 51 | 11.19 | 6 | -60 | -57 | R Cerebelum (IX) |
|  | 119 | 10.01 | -15 | -30 | 66 | L Paracentral Lobule |
|  |  | 9.00 | -6 | -21 | 69 | L Paracentral Lobule |
|  |  | 8.00 | 33 | -18 | 69 | R Precentral Gyrus |

L=left; R=right; IFG=inferior frontal gyrus; SLN=semantic language network; ECN=executive control network; HVN=higher visual network; PVN=primary visual network; DMN=default-mode network.
